# Population dynamics of GC-changing mutations in humans and great apes

**DOI:** 10.1101/2020.09.25.313411

**Authors:** Juraj Bergman, Mikkel Heide Schierup

## Abstract

**Background:** The nucleotide composition of the genome is a balance between origin and fixation rates of different mutations. For example, it is well-known that transitions occur more frequently than transversions, particularly at CpG sites. Differences in fixation rates of mutation types are less explored. Specifically, recombination-associated GC-biased gene conversion (gBGC) may differentially impact GC-changing mutations, due to differences in their genomic distributions and efficiency of mismatch repair mechanisms. Given that recombination evolves rapidly across species, we explore gBGC of different mutation types across human populations and among great ape species.

**Results:** We report a stronger correlation between GC frequency and recombination for transitions than for transversions. Notably, CpG transitions are most strongly affected by gBGC. We show that the strength of gBGC differs for transitions and transversions but that its overall strength is positively correlated with effective population sizes of human populations and great ape species, with some notable exceptions, such as a stronger effect of gBGC on non-CpG transitions in populations of European descent. We study the dependence of gBGC dynamics on flanking nucleotides and show that some mutation types evolve in opposition to the gBGC expectation, likely due to hypermutability of specific nucleotide contexts.

**Conclusions:** Differences in GC-biased gene conversion are evident between different mutation types, and dependent on sex-specific recombination, population size and flanking nucleotide context. Our results therefore highlight the importance of different gBGC dynamics experienced by GC-changing mutations and their impact on nucleotide composition evolution.

## 1 Introduction

The nucleotide composition of the genome evolves under a composite of evolutionary forces, often acting antagonistically. Mutations that are introduced into the genome are generally AT-biased, *i.e*., the mutation rate from strong (GC) to weak (AT) nucleotides is higher than in the opposite direction. However, the proportion of fixed sites in the genome is GC-biased compared to the expectation based on mutation rate differences. This observation is usually ascribed to a biased, and as of yet unidentified, recombination-associated mismatch repair mechanism, termed GC-biased gene conversion; gBGC [14, 17, 25]. Ample evidence from mammals – mainly the repeatedly observed positive correlation between recombination rate and GC content [15, 16, 34, 35, 41, 43, 57] – points to gBGC as a major determinant of nucleotide composition at the genomic level.

The proportions of different mutation types, *i.e*., the mutational spectrum, is known to vary with the surrounding nucleotide context, as well as to evolve rapidly at the species and population levels in humans and great apes [8, 22, 23, 30]. However, the fixation biases of different mutation types are not nearly as well-characterised, even at the basic level of transition and transversion differences. With this study, we aim to provide a broader perspective on mutation fixation biases, particularly with respect to potential differences in gBGC dynamics experienced by different GC-changing mutations. We explore multiple possible sources of these differences, including mutation-specific repair mechanisms, differences in genomic distributions of mutations, surrounding genomic context as well as differences in the effective sizes and recombination rates between sexes, populations and species.

During meiotic recombination, segregating variants can come into contact in recombination heteroduplexes which consist of homologous pairings of DNA strands between chromosomes of different parental origins. Homology search, as well as mismatch repair (MMR) required for proper heteroduplex formation share much of the molecular machinery of post-replicative MMR [9, 26, 51]. However, while MMR that occurs immediately after replication utilizes epigenetic marks to distinguish between the template and nascent DNA strands and restore the mismatched base pair, the repair of mismatches in recombination heteroduplexes is context-dependent. Specifically, heteroduplex mismatches underlyed by GC-changing mutations are more often resolved into G:C than A:T pairs, providing the basis for gBGC. GC-changing mutations that precede gBGC are either G(C)↔A(T) transitions or G(C)↔T(A) GC-changing transversions. Therefore, a heteroduplex mismatch that forms during meiotic recombination will be of the G:T or C:A type if a site segregates for a transition, or of the G:A or C:T type for transversion-segregating sites (Supplementary figure 1). Molecular studies of *Escherichia coli* MMR efficiency indicate that different mismatch types are successfully resolved at different rates – generally, G:T and C:A mismatches are better substrates for the MMR machinery than G:A or C:T mismatches [13, 28, 54]. Studies on human cell cultures yielded similar results [24, 56]. Additionally, biased repair of heteroduplex mismatches into G:C, rather than A:T pairs, has been reported in various mammalian systems, especially for transition-associated mismatches [4, 6, 60, 61]. Together, these results describe molecular underpinnings that may potentially manifest as different gBGC dynamics of transitions and transversions at the population genomic level.

The genomic distribution of different mutations can further modify the gBGC effect experienced by different mutation types. Specifically, as the transition/transversion ratio of new mutations varies along the genome [48], the proportion of transitions in high (or low) recombination regions can be different than the same proportion for transversions. As gBGC is driven by recombination rate, this inequality could result in different average gBGC strength experienced by the two mutation types. Most notably, the particularly high proportion of CpG transitions in GC-rich regions with high recombination rates [44, 47], are likely to experience the strongest gBGC dynamics compared to other mutation types.

The strength of gBGC is usually quantified as *B* = *N_e_b*, where *N_e_* is the effective population size and *b* is the conversion bias parameter defined by [37]. Given this parametrization, nucleotide diversity is affected in a manner indistinguishable from directional selection towards GC content. The *B* parameter can be calculated from the skew of the allele frequency spectrum (AFS) – a categorical representation of nucleotide diversity within a sample of individuals from a particular population. AFS-based methods have been recently applied to both human [18] and great ape datasets [5]. [18] considered a model where the ancestral state of segregating variants is assumed to be known (*i.e*., polarized data), making it possible to contrast the AFS of derived weak (W = AT) to strong (S = GC) mutations against the S→W AFS. On the other hand, [5] used unpolarized data and considered AFSs of all six possible nucleotide pairs to estimate GC bias. Both studies yielded gBGC estimates of *B* < 1. Even though these estimates are in the nearly neutral range of fixation coefficients [39, 40], they nonetheless have a profound effect on the evolution of nucleotide content as millions of sites across the genome are potentially affected by gBGC.

Here, we use human [10, 27] and great ape population genomic data sets [12, 38, 42, 63] to assess gBGC dynamics of the three most common mutation types: CpG and non-CpG transitions, and GC-changing transversions. We relate patterns of nucleotide variation to population-specific recombination maps that have been recently derived for humans [21, 50], or that we reconstruct *de novo* within this study for eight subspecies of great apes. We find a stronger correlation between the recombination rate and segregating GC frequency for transitions compared to GC-changing transversions. We estimate the GC fixation bias and observe that CpG transitions have an especially large *B* value compared to the other mutation types, likely due to the strong correlation between their genomic distribution and recombination rate. The observed patterns are largely congruent with differences in historical effective sizes across populations and species.

## 2 Results

### 2.1 GC-changing mutations and recombination rate

As GC-biased gene conversion (gBGC) only affects GC-changing mutations during recombination, we focus our analyses on population dynamics of transitions and G(C)↔T(A) transversions. The basis of all our analyses is the GC frequency of a segregating GC-changing mutation that we calculate for each bi-allelic site as the proportion of haploid genomes, in a respective population sample, that have a G(C) as opposed to an A(T) at a transition-segregating site, or a G(C) *v.s*. a T(A) at a transversion segregating site. For most of the analyses of human data we focus on three populations from the 1000 Genomes Project [10] – African Yoruba; YRI, Italian Toscana; TSI and Han Chinese; CBH – as representatives of the ancestral and initial Out-of-Africa human populations. The Icelandic (ISL) population was also studied as nucleotide diversity and the recombination map of this population has the highest resolution of any human population studied to date.

In accordance with a genome-wide effect of gBGC dynamics, we observe a positive correlation between the average recombination rate and average segregating GC frequency of GC-changing mutations in 1 Mb genomic windows of the African YRI population (Spearman’s *ρ* = 0.3757, *p* < 2.2×10^−16^; Figure 1A). This pattern is also consistent across Eurasian populations (Table 1). We next separate GC-changing mutations into transitions and transversion to investigate potential differences between their population dynamics with respect to recombination-associated processes. We distinguish between CpG and non-CpG transitions, as a significant proportion of GC-changing mutations are transitions at CpG sites (27.78%) that likely experience distinct population dynamics due to high recombination rates in GC-rich genomic regions [15]. Of the remainder of GC-changing mutations, non-CpG transitions are the most abundant mutation type (53.13%), while non-CpG and CpG transversions comprise 15.94% and 3.14%, respectively. These proportions are based on the African YRI population. We will omit CpG transversions from this part of the analysis and revisit their particular dynamics in the section regarding flanking context dependency.

**Figure 1:**
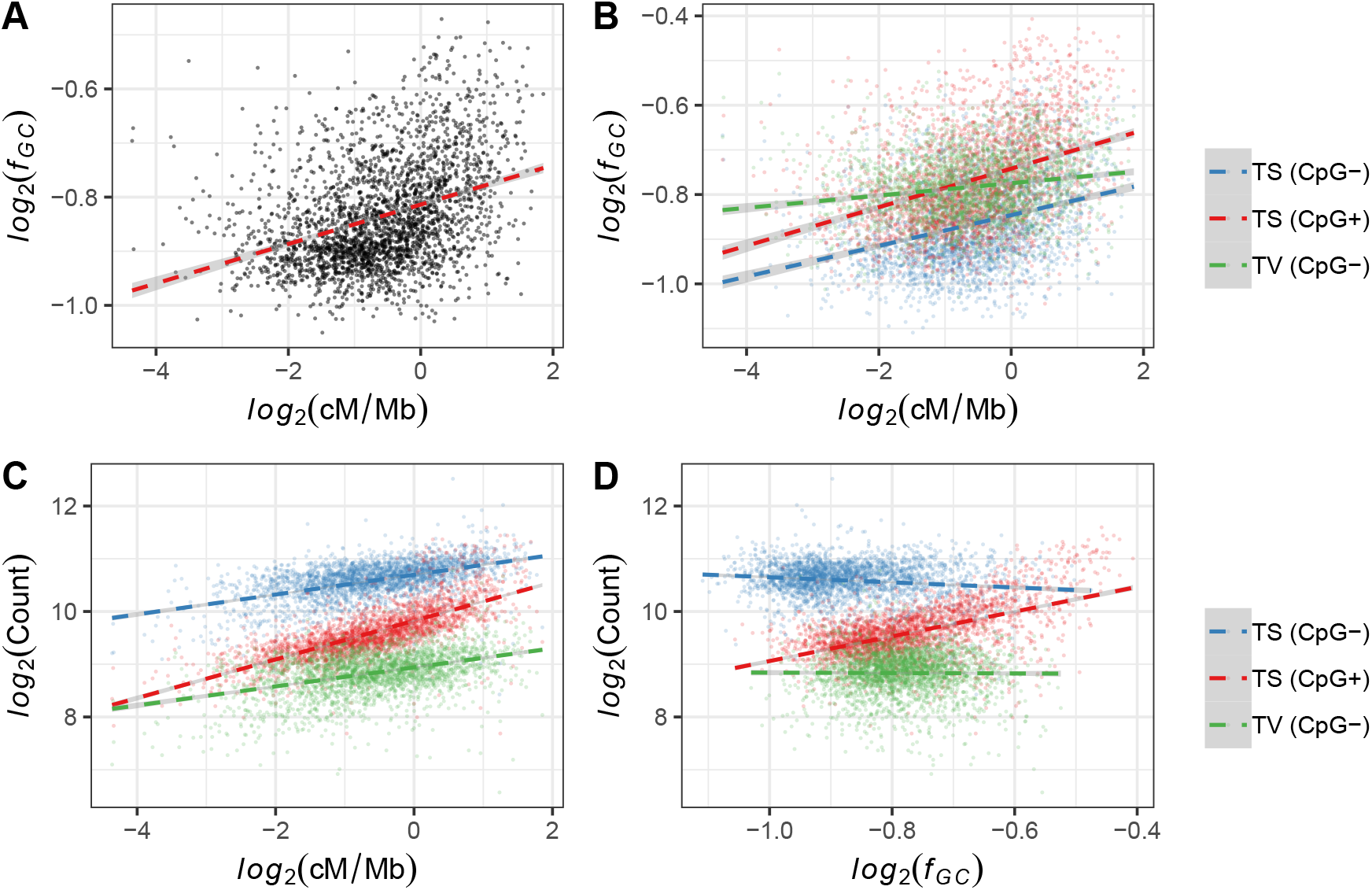
Relationship between the average GC frequency (*f*_GC_) at segregating sites and average recombination rate in 1 Mb autosomal windows of the African Yoruba population, for **A.** all GC-changing mutations, and **B.** for non-CpG transitions (TS; CpG−), CpG transitions (TS; CpG+) and GC-changing transversions (TV; CpG−) separately. **C.** Relationship between the count of GC-changing segregating sites and recombination rate in 1 Mb autosomal windows of the African Yoruba population for non-CpG transitions (TS; CpG−), CpG transitions (TS; CpG+) and GC-changing transversions (TV; CpG−) separately. **D.** Relationship between the count of GC-changing segregating sites and average GC frequency (*f*_GC_) in 1 Mb autosomal windows of the African Yoruba population for non-CpG transitions (TS; CpG−), CpG transitions (TS; CpG+) and GC-changing transversions (TV; CpG−) separately.

**Table 1:** Linear regression coefficients with corresponding *R*^2^ values for the relationship between the average GC frequency (*f*_GC_) or count proportion of all segregating sites, non-CpG transitions (TS; CpG−), CpG transitions (TS; CpG+) or GC-changing transversions (TV; CpG−) and average recombination rate across 1 Mb autosomal regions of different human populations (YRI = Yoruba; TSI = Toscana; CHB = Han Chinese; ISL = Iceland). All coefficients are significant at *p* < 0.0001.

| Parameter | Population |  |  |  |
| --- | --- | --- | --- | --- |
|  | YRI | TSI | CHB | ISL |
| $f_{GC}$ (All) | 0.0294 ($R^2 = 0.1608$) | 0.0201 ($R^2 = 0.1437$) | 0.0154 ($R^2 = 0.1393$) | 0.0154 ($R^2 = 0.1464$) |
| $f_{GC}$ (TS; CpG-) | 0.0267 ($R^2 = 0.1303$) | 0.0184 ($R^2 = 0.1111$) | 0.0137 ($R^2 = 0.1007$) | 0.0141 ($R^2 = 0.1166$) |
| $f_{GC}$ (TS; CpG+) | 0.0363 ($R^2 = 0.1752$) | 0.0253 ($R^2 = 0.1477$) | 0.0194 ($R^2 = 0.1401$) | 0.0188 ($R^2 = 0.1551$) |
| $f_{GC}$ (TV; CpG-) | 0.0119 ($R^2 = 0.0420$) | 0.0068 ($R^2 = 0.0180$) | 0.0048 ($R^2 = 0.0133$) | 0.0092 ($R^2 = 0.0567$) |
| %Count (All) | 0.0111 ($R^2 = 0.3776$) | 0.0112 ($R^2 = 0.3580$) | 0.0091 ($R^2 = 0.3086$) | 0.0056 ($R^2 = 0.2640$) |
| %Count (TS; CpG-) | 0.0077 ($R^2 = 0.2149$) | 0.0081 ($R^2 = 0.2105$) | 0.0066 ($R^2 = 0.1715$) | 0.0039 ($R^2 = 0.1429$) |
| %Count (TS; CpG+) | 0.0189 ($R^2 = 0.4695$) | 0.0184 ($R^2 = 0.4427$) | 0.0149 ($R^2 = 0.3991$) | 0.0100 ($R^2 = 0.3721$) |
| %Count (TV; CpG-) | 0.0076 ($R^2 = 0.1468$) | 0.0081 ($R^2 = 0.1512$) | 0.0064 ($R^2 = 0.1240$) | 0.0036 ($R^2 = 0.0852$) |

As expected, the correlation between average segregating GC frequency and recombination rate in 1 Mb genomic windows for different mutation types is positive and significant (Spearman’s *ρ* = 0.3480, *p* < 2.2×10^−16^ for non-CpG transitions; Spearman’s *ρ* = 0.3924, *p* < 2.2×10^−16^ for CpG transitions; Spearman’s *ρ* = 0.1753, *p* < 2.2×10^−16^ for transversions; Figure 1B). We further observe that this correlation is stronger for transitions compared to the same correlation for transversions (*t* = 12.0153, *p* < 0.001 for the comparison between CpG transitions and transversions; *t* = 10.8384, *p* < 0.001 for the comparison between non-CpG transitions and transversions). Notably, the GC frequency distributions differ between the three mutation types both with respect to their means and variances. The mean of GC frequencies across 1 Mb regions is higher for CpG transitions compared to non-CpG transversions (*t* = 6.528, *p* = 7.364× 10^−11^), which have a higher mean compared to non-CpG transitions (*t* = 33.339, *p* < 2.2×10^−16^). The differences between the distribution means are likely the consequence of different gBGC dynamics experienced by these mutation types. Furthermore, average GC frequencies of transitions are more variable compared to transversions (Levene test statistic = 107.72, *p* < 2.2^−16^ for the comparison between CpG transitions and non-CpG transversions; Levene test statistic = 300.86, *p* < 2.2^−16^ for the comparison between non-CpG transitions and non-CpG transversion), and GC frequencies of CpG transitions are more variable compared to non-CpG transitions (Levene test statistic = 50.45, *p* = 1.382^−12^). Therefore, the differences between correlations in Figure 1B are likely driven by the wider range of average GC frequencies for transitions compared to transversions. Linear regression analysis of the YRI population (Table 1) predicts that an increase of 2.67% and 3.63% in average GC frequency is expected for non-CpG and CpG transitions, respectively, per 1 cM/Mb increase in recombination rate, while only a 1.19% increase is expected for transversions. The recombination rate explains ~ 13 – 18% of variance in average GC frequency for transition sites, and ~ 4% for transversions. The expected increase in average GC frequency per unit of recombination rate for the Eurasian populations is consistent with these results, but lower compared to the YRI population (Table 1). This observation is in accordance with the historically lower effective sizes of these populations, despite the fact that a higher recombination rate is inferred for Eurasian populations (Supplementary figure 2A). This observation indicates that the effective size, rather than differences between recombination maps, is a more important determining factor of between-population differences in gBGC strength. Generally, correlations and regression coefficients are more similar between all sites and transitions, rather than transversions, due to the fact that the ratio of segregating transitions and GC-changing transversions is biased – approximately 4:1 in favour of transitions. Therefore, the genome-wide profile of gBGC dynamics is mainly due to transition-specific processes.

The correlation between the abundance of segregating sites and recombination rate is positive (Spearman’s *ρ* = 0.5089, *p* < 2.2×10^−16^ for non-CpG transitions; Spearman’s *ρ* = 0.7689, *p* < 2.2×10^−16^ for CpG transitions; Spearman’s *ρ* = 0.3762, *p* < 2.2×10^−16^ for transversions; Figure 1C), indicating that mutations generally accumulate in high recombination regions. Furthermore, this correlation is strongest for CpG transitions compared to the two other mutation types (*t* = 25.0652, *p* < 0.001 for the comparison between CpG and non-CpG transitions; *t* = 31.7503, *p* < 0.001 for the comparison between CpG transitions and non-CpG transversions). We next quantify the effect of recombination on mutation abundance for each mutation type by calculating linear regression coefficients for the relationship between the count proportion of a specific mutation type (with respect to the corresponding total count) and average recombination rate in 1 Mb windows (Table 1). We opt to use count proportions, rather than the absolute counts of mutations, to account for the differences between the total numbers of different mutation types. We find that the recombination rate explains ~ 37-47% of variance in the proportion of CpG transitions, compared to ~ 14-21% and ~ 9-15% for non-CpG transitions and transversions, respectively (Table 1). For the YRI population, an increase in one unit of recombination rate results in an increase of 0.0189% of total CpG transitions, compared to 0.0077% and 0.0076% for non-CpG transitions and transversions, respectively. A similar trend is true for Eurasian population. These results demonstrate that CpG transitions are especially enriched in high-recombination regions and are expected to evolve under the strongest gBGC dynamics. Notably, the ISL population has a considerably lower increase in count proportion per unit increase of recombination when compared to other Eurasian populations. This is likely due to the different inference methodology and higher resolution of the ISL recombination map, resulting in a larger variance of recombination rates (Supplementary figure 2A).

As the average GC frequency and abundance of segregating mutations is positively correlated to recombination rate (Figure 1A-C), we may assume that recombination-associated mutations contribute to the observed population dynamics of GC-changing mutations. However, although there is indeed a higher mutation rate in high-recombination regions, putative recombination-associated mutations comprise a very small fraction of genome-wide *de novo* mutations (DNMs) – in a recent study, only 173 out of 200,435 DNMs were found within 1 kb from a recombination crossover event [21]. Notably, an increase in mutation rate close to maternal crossover events can extend up to 40 kb; however, these mutations are mostly C→G transversions that are unaffected by gBGC. Furthermore, direct sequencing of recombination hotspots in human sperm cells demonstrates that, despite a higher mutation rate in these regions, the opposing effect of gBGC is nonetheless the dominant factor in determining the sequence evolution of high-recombination regions [1].

The analyses above were done using sex-averaged recombination rates. However, as sex-specific recombination maps are available for the ISL population, this allows us to specifically asses the influence of sex on the population dynamics of GC-changing mutations. Namely, we repeat the linear regression analysis from Table 1 with a model where the explanatory variables include male and female recombination rates, as well as their interaction (Table 2). Notably, the regression coefficients are positive and significant for both the maternal and paternal recombination maps, signifying that recombination in both sexes is associated with the increase in average GC frequency and abundance of mutations. However, the magnitude of the coefficients is always higher for the paternal recombination rate, suggesting that the dynamics of GC-changing mutations are more strongly influenced by paternal recombination, despite the fact that the genome-wide maternal recombination rate is higher (Supplementary figure 2A). Interestingly, the interaction coefficient is always negative, implying that maternal and paternal recombination have antagonistic effects. These observations are likely a consequence of the differences between male and female meiosis.

**Table 2:** Linear regression coefficients with corresponding *R*^2^ values for the relationship between the average GC frequency (*f*_GC_) or count proportion of all segregating sites, non-CpG transitions (TS; CpG−), CpG transitions (TS; CpG+) or GC-changing transversions (TV; CpG−) and recombination rate parameters across 1 Mb autosomal regions for the ISL population, where *r_m_* is the maternal recombination rate and *r_p_* is the paternal recombination rate.

| Parameter | Explanatory variables | | | $R^2$ |
| --- | --- | --- | --- | --- |
| | $r_m$ | $r_p$ | $r_m \times r_p$ | |
| $f_{GC}$ (All) | 0.0105*** | 0.0186*** | -0.0044*** | 0.147 |
| $f_{GC}$ (TS; CpG-) | 0.0113*** | 0.0173*** | -0.0048*** | 0.119 |
| $f_{GC}$ (TS; CpG+) | 0.0112*** | 0.0232*** | -0.0050*** | 0.1582 |
| $f_{GC}$ (TV; CpG-) | 0.0070*** | 0.0078*** | -0.0019** | 0.0488 |
| %Count (All) | 0.0023*** | 0.0061*** | -0.0008*** | 0.2776 |
| %Count (TS; CpG-) | 0.0017*** | 0.0035*** | -0.0003* | 0.1408 |
| %Count (TS; CpG+) | 0.0042*** | 0.0123*** | -0.0020*** | 0.4345 |
| %Count (TV; CpG-) | 0.0009** | 0.0036*** | -0.0001 (*n.s.*) | 0.0903 |

\*\*\* $p < 0.001$ ; \*\* $p < 0.01$ ; \* $p < 0.05$ ; *n.s.* = non-significant

### 2.2 Distribution of GC frequencies

The distribution of GC frequencies for transition- and transversion-segregating sites has the characteristic U-shape, typical for human populations with a relatively low population size-scaled mutation rate *θ* (4*N_e_μ*) < 0.01 (Figure 2). All distributions are skewed towards high GC frequencies, as expected for sites evolving under gBGC dynamics. For the YRI population, the distribution of GC frequencies for CpG transitions has a more extreme skew compared to both non-CpG transition (*t* = 155.81, *p* < 2.2×10^−16^) and transversion distributions (*t* = 40.973, *p* < 2.2×10^−16^), with a higher mean (60.01% GC for CpG transitions, 54.90% GC for non-CpG transitions and 58.21% GC for transversions) and median (79.17% GC for CpG transitions, 67.59% GC for non-CpG transitions and 75.93% GC for transversions; Supplementary table 1). A similar trend is observed for Eurasian populations, but with a general reduction in proportions of low and high GC frequency variants likely due to the bottleneck-induced reduction of effective sizes in these populations following the Out-of-Africa migration. The skewness of the GC frequency distribution reflects the strength of gBGC dynamics acting on each mutation type and is in accordance with the observed correlations in Figure 1B, C. Additionally, these distributions further indicate that CpG transitions evolve under strongest gBGC dynamics across all human populations.

**Figure 2:**
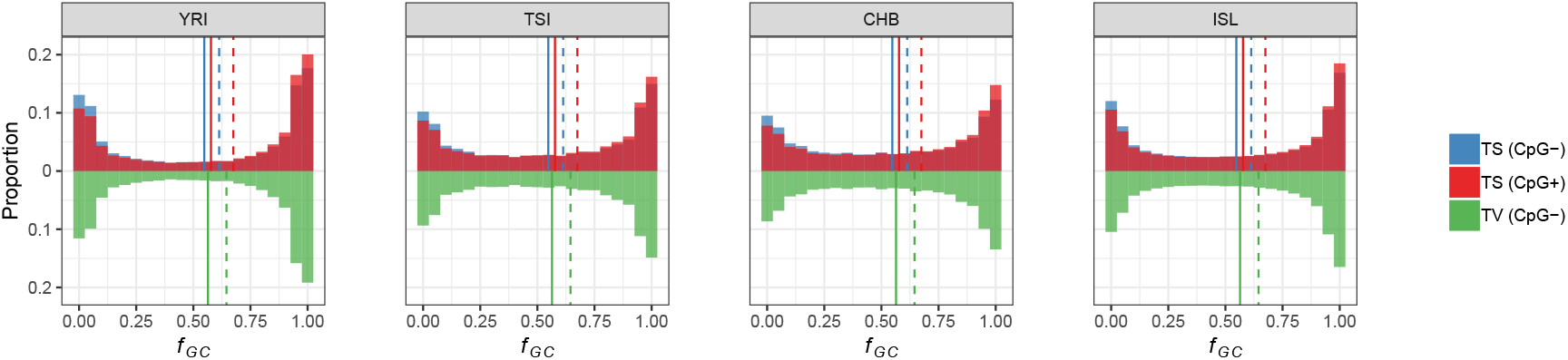
Distribution of GC frequencies (*f*_GC_) for non-CpG transitions (TS; CpG−), CpG transitions (TS; CpG+) and GC-changing transversions (TV; CpG−) of Yoruba (YRI), Toscana (TSI), Han Chinese (CHB) and Iceland (ISL) populations. The solid lines represent the medians, and the dashed lines represent the means of the distributions.

The distribution of GC frequencies differs between non-CpG transitions and transversions (Figure 2), with transversions having a significantly higher average GC frequency (*t* = 82.452, *p* < 2.2×10^−16^), despite a stronger correlation between recombination rate and average GC frequency for non-CpG transitions (Figure 1B). A contributing factor to this observation is the fact that the number of non-CpG transitions is negatively correlated with their average GC frequency (Spearman’s *ρ* = −0.0782, *p* < 0.0001), while the same correlation is positive for CpG transitions (Spearman’s *ρ* = 0.4841, *p* < 2.2×10^−16^) and transversions (Spearman’s *ρ* = 0.0396, *p* = 0.0408; Figure 1D). In other words, the proportion of non-CpG transitions in high-vs. low-recombination regions is relatively smaller than the corresponding proportion of transversions, thus contributing to their less skewed GC frequency distribution. Another likely contributor is the higher variance of GC frequencies for non-CpG transitions compared to transversions (Figure 1B).

The skew in GC frequency distributions, such as the one observed in Figure 2, could be a consequence of the difference in mutation rates between GC→AT and AT→GC mutations. However, in the case of low mutation rates *θ*, as are typical for humans and great apes, the skewness of the GC frequency distribution of segregating sites should be only slightly influenced by mutation processes and largely determined by the fixation bias towards the preferred allele [32, 58]. The strongest potential impact of mutation on the GC frequency skew should be present at CpG sites, as CpG→TpG transitions have a ~ 10 × higher per base mutation rate compared to the genome-wide rate [21]. However, Supplementary figure 3 shows that the mutation rate for human CpG transitions is capable of introducing only a very minor skew in the distribution of GC frequencies, while a 10-100 × higher CpG→TpG mutation rate than the one estimated would be required to account for the skew of the GC frequency distributions in Figure 2.

### 2.3 Quantification of the fixation bias in human populations

To quantify the effect of gBGC on segregating sites, we use allele frequency spectra (AFSs) that are constructed by assigning sites into 50 equally-sized categories based on their segregating GC frequency. Such AFSs have been successfully used to assess fixation biases [5, 58], without requiring the knowledge of the ancestral state of the segregating site. The fixation bias *B* is parametrized as the product of the effective population size *N_e_* and the conversion bias *b* [37]. For the YRI population, average *B* estimates calculated using all autosomal sites are 0.3072, 0.6362 and 0.5188 for non-CpG transitions, CpG transitions and non-CpG transversions, respectively. These estimates are in the nearly neutral range of selection coefficients [39, 40] and in line with previous estimates for humans [18]. Consistent with this result, the per chromosome *B* estimates (Figure 3A) are higher for CpG transitions than non-CpG transitions (*t* = 7.3349, *p* < 0.001) or transversions (*t* = 3.7355, *p* < 0.001). The *B* estimates for CpG transitions are approximately 2.07× and 1.23× larger compared to non-CpG transitions and transversions, respectively. These observations are reflected in the strong correlation between the average segregating GC frequency of CpG transitions and recombination rate (Figure 1B) and their accumulation in high-recombination regions (Figure 1C).

**Figure 3:**
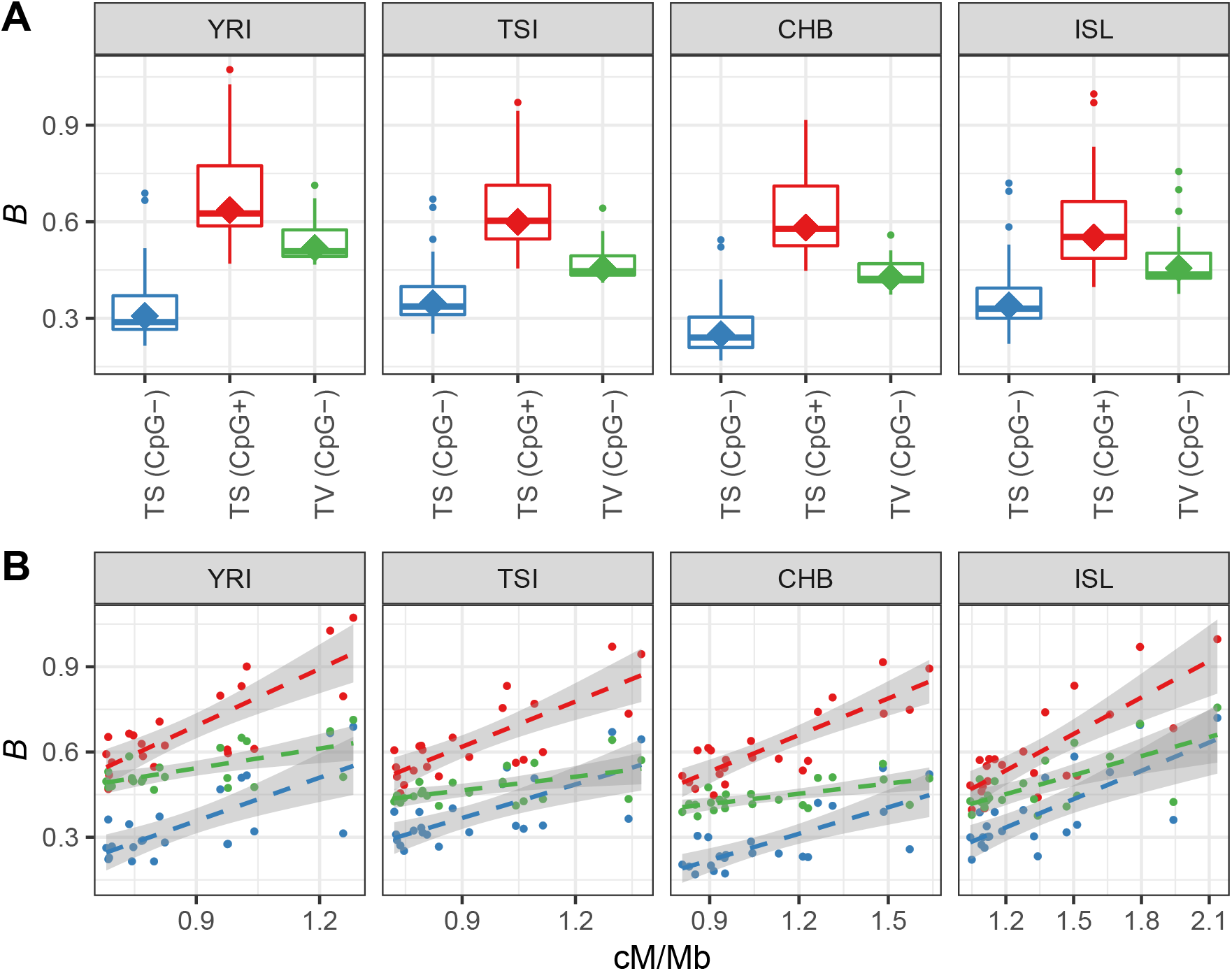
Distribution of the per-chromosome *B* estimates for non-CpG transitions (TS; CpG−), CpG transitions (TS; CpG+) and GC-changing transversions (TV; CpG−). The “diamond” points correspond to the genome-wide *B* estimated for all autosomes together. **B.** Relationship between the non-CpG transitions (blue), CpG transitions (red) or GC-changing transversions (green) *B* parameter and the sex-averaged recombination rate across autosomes. Each point is an autosomal chromosome and the four panels in each figure correspond to the Yoruba (YRI), Toscana (ISL), Han Chinese (CHB) and Iceland (ISL) populations.

Estimates of *B* for CpG transitions and non-CpG transversions are in correspondence with the differences in effective sizes of populations, with YRI having the highest *B* estimates compared to the Eurasian populations. However, the difference is only significant when comparing estimates of non-CpG transversions between the YRI population and the Eurasian populations (*t* = 3.3516, *p* = 0.002 for the YRI-TSI comparison; *t* = 5.1994, *p* < 0.001 for the YRI-CHB comparison; *t* = 2.0622, *p* = 0.0461 for the YRI-ISL comparison). The *B* estimates for non-CpG transitions are lowest in the CHB population consistent with the lower effective size, but not significant in comparison with the other populations. Another notable trend is the higher *B* estimates of non-CpG transitions in European populations compared to Africans and Asians. This trend is in opposition to the historical difference in population sizes between the African and European populations. It is also unlikely to be a product of the generally higher recombination rate in European populations, as we would then expect a similar effect on the two other mutation types. Furthermore, the correlations between the African YRI and European TSI populations for the GC frequency (Spearman’s *ρ* = 0.8799, *p* < 2.2×10^−16^; Supplementary figure 4A) and abundance (Spearman’s *ρ* = 0.8679, *p* < 2.2×10^−16^; Supplementary figure 4B) of non-CpG transitions is highly significant at the 1 Mb scale, as is the correlation of recombination rates (Spearman’s *ρ* = 0.9268, *p* < 2.2×10^−16^; Supplementary figure 4C). Therefore, this trend may be a consequence of genomic features that are only detectable at a finer genomic scale, such as the difference in location and intensity of recombination hotspots between African and European populations. Interestingly, the observed trends from Figure 3 are also conserved at the geographic superpopulation level (Figure 4 and Supplementary figure 5), suggesting that gBGC processes have been stable after population establishment following the Out-of-Africa migration.

**Figure 4:**
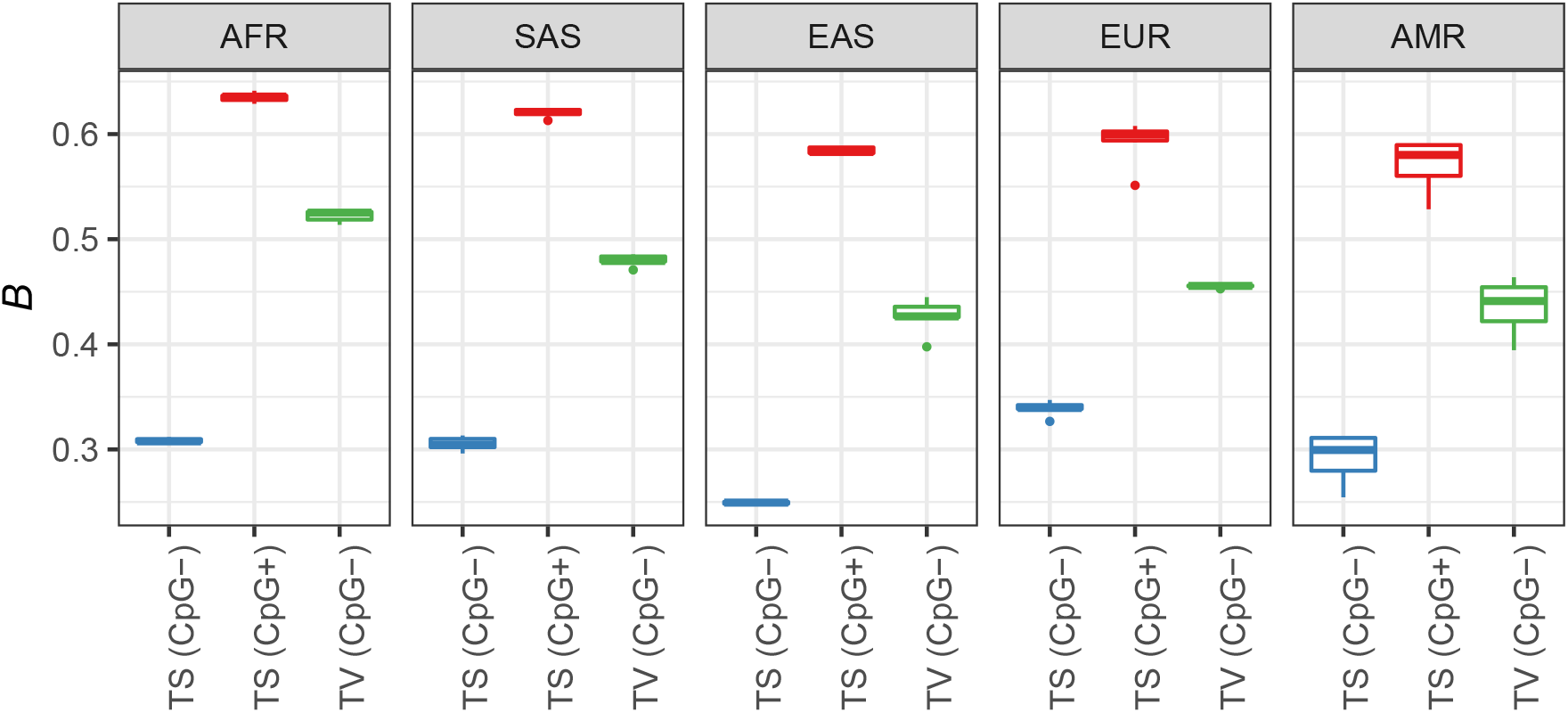
Distribution of the genome-wide *B* estimates for non-CpG transitions (TS; CpG−), CpG transitions (TS; CpG+) and GC-changing transversions (TV; CpG−) across populations within different geographic superpopulations (AFR = African; SAS = South Asian; EAS = East Asian; EUR = European; AMR = Admixed American).

We next investigate the relationship between *B* and recombination rate – as expected, for all populations and mutation types, there is a positive correlation between *B* and the average recombination rate at the chromosome level (Figure 3B). Chromosome-level recombination rates explain a substantial proportion of the variance in per chromosome *B* estimates (Table 3) - on average ~ 47-53% for non-CpG transitions, ~ 60-69% for CpG transitions and ~ 23-42% for non-CpG transversions. Furthermore, the YRI recombination rate varies at a similar scale to Eurasian populations (Figure 3B), yet has the highest increase in *B* per increase in unit of recombination (Table 3). Again, this observation demonstrates that the efficiency of gBGC is promoted by the historically higher effective size of the African population.

**Table 3:** Linear regression coefficients with corresponding *R*^2^ values for the relationship between per-chromosome *B* of non-CpG transitions (TS; CpG−), CpG transitions (TS; CpG+) or GC-changing transversions (TV; CpG−) and recombination rate, for different human populations (YRI = Yoruba; TSI = Toscana; CHB = Han Chinese; ISL = Iceland).

| Parameter | Population |  |  |  |
| --- | --- | --- | --- | --- |
|  | YRI | TSI | CHB | ISL |
| $B(\text{TS; CpG}^-)$ | 0.5072 ($R^2 = 0.4894$)*** | 0.3942 ($R^2 = 0.4665$)*** | 0.3106 ($R^2 = 0.5294$)*** | 0.3322 ($R^2 = 0.5170$)*** |
| $B(\text{TS; CpG}^+)$ | 0.6673 ($R^2 = 0.6268$)*** | 0.5302 ($R^2 = 0.5981$)*** | 0.4286 ($R^2 = 0.6905$)*** | 0.4280 ($R^2 = 0.6113$)*** |
| $B(\text{TV; CpG}^-)$ | 0.2302 ($R^2 = 0.3462$)** | 0.1597 ($R^2 = 0.2683$)** | 0.1199 ($R^2 = 0.3759$)** | 0.2233 ($R^2 = 0.4219$)*** |

\*\*\* $p < 0.001$ ; \*\* $p < 0.01$

### 2.4 GC-changing mutations and flanking nucleotide context

In this section we classify GC-changing mutations and their reverse complements into 32 mutational types depending on their flanking 5’ and 3’ nucleotides. We note that the proportions of different mutation types vary widely, with the 5’ACG3’↔5’ATG3’ type being the most abundant (Figure 5A). Generally, transitions, and especially CpG transitions, are the most abundant mutation types. Furthermore, the proportions of specific mutation types are very similar between populations. However, we do recover the strong enrichment signal of the 5’TCC3’↔5’TTC3’ mutation type in European populations, compared to the YRI and CHB populations (Supplementary figure 6), as previously observed [23].

**Figure 5:**
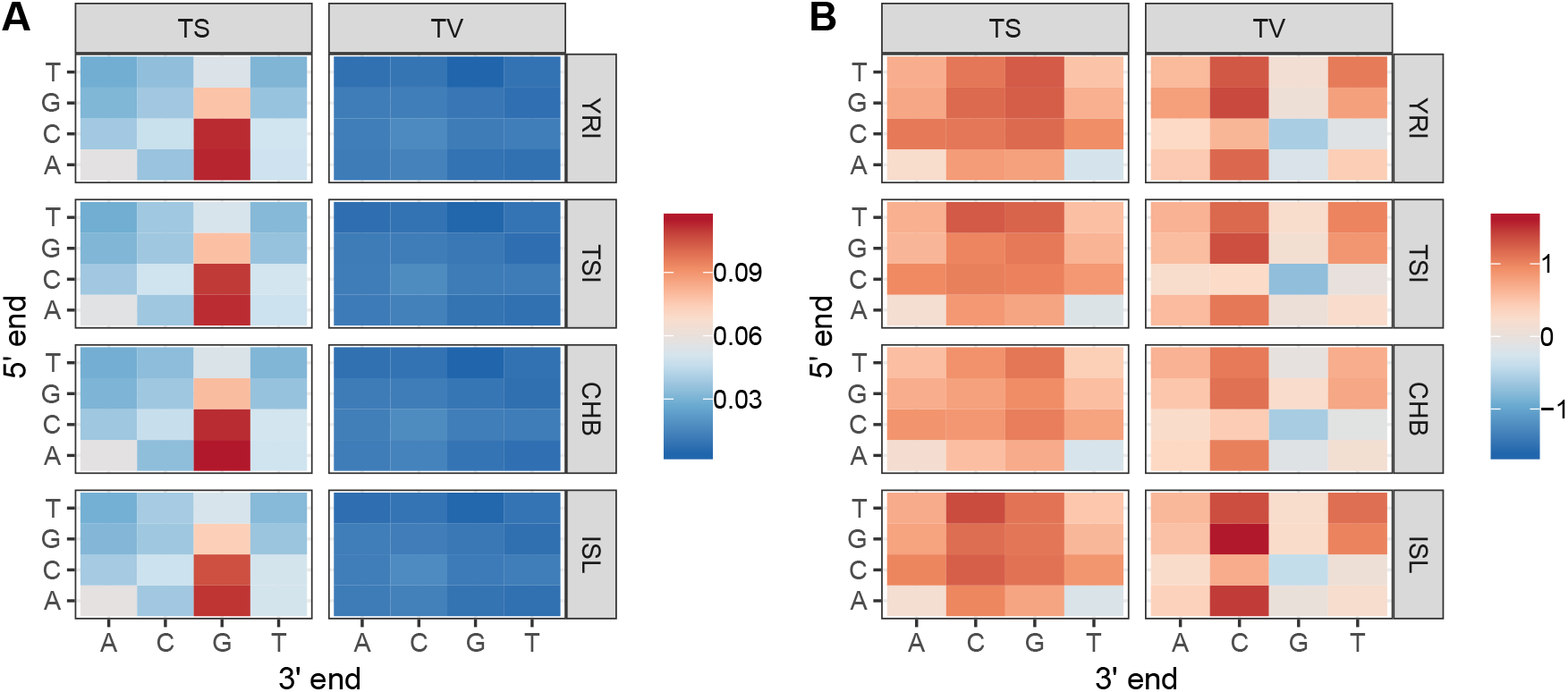
**A.** The proportions of the 32 different mutation types for the YRI, TSI, CHB and ISL populations. **B.** The values of the fixation bias *B* of 32 different mutation types for the the YRI, TSI, CHB and ISL populations.

We next calculate the fixation index *B* for each of the 32 mutation types (Figure 5B). As expected, most *B* values are positive, indicating fixation towards GC content as expected for sites evolving under GC-biased gene conversion. Interestingly, the highest *B* values are estimated for 5’*CC3’↔5’*AC3’ transversions, demonstrating that subsets of non-CpG transversions may experience gBGC dynamics stronger than even CpG transitions. Unexpectedly however, some mutation types have negative *B* values. For transitions, the 5’ACT3’↔5’ATT3’ mutation type has a negative *B* value in all populations, in opposition to the gBGC expectation. For transversions, negative *B* values are present for the 5’CCT3’↔5’CAT3’ mutation type and two types of CpG transversions (5’CCT3’↔5’CAT3’ and 5’ACG3’↔5’AAG3’). This unexpected result is likely due to non-gBGC processes affecting the GC frequency distribution of these mutation types. We next focus on CpG transversions, which seem to be the most affected by this phenomenon.

Transversions that segregate at CpG sites have the following characteristics. Firstly, they are the most rare mutation type of all GC-changing mutations (comprising 3.14%; 250,334 sites in total; Supplementary table 1). Secondly, their average segregating GC frequency is lowest out of all the mutation types with a mean of 40.27% GC and median of 21.76% GC (Supplementary table 1). Accordingly, the AFS of CpG transversions is skewed towards low GC (*i.e*., high AT) frequency variants (Supplementary figure 7). However, despite their low GC content, CpG transversions have typical characteristics of sites evolving under gBGC dynamics, such as the positive correlation between average GC frequency and recombination (Supplementary figure 8A). Furthermore, the count of CpG transversions is positively correlated to recombination rate, as well as average GC frequency (Supplementary figure 8B, C). Therefore, the GC frequency distribution of CpG transversions is affected either by an excess of low GC variants or conversely, the lack of high GC variants. An excess of low GC variants would imply an exceptionally high AT→GC mutation rate, which is highly unlikely given that mutation rates in humans are generally low and AT-biased. Furthermore, under this scenario, we would expect a much higher representation of CpG transversions, as opposed to only 3.14% of all GC-changing mutations. Another possible explanation for the skew in the CpG transversion AFS is a biased fixation of AT nucleotides at CpG transversion sites. However, this is also highly unlikely, as no biological mechanisms are known that would cause such a bias. Therefore, a more likely explanation for the observed GC frequency distribution of CpG transversions is a lack of high frequency GC variants. This underrepresentation can occur due to the high mutability of GC content at CpG sites. Specifically, as sites that segregate for CpG transversions reach high GC frequency, the probability of additional mutations increases, in turn decreasing the number of detectable CpG transversions segregating at high GC frequency. As a result, high GC frequency CpG transversions would be enriched for tri-nucleotide variants, harder to detect with conventional SNP calling methods and possibly mis-classified into other mutation types, resulting in their general under-representation. Additionally, recent shifts in the recombination map can also contribute to the observed patterns. While it is difficult to reliably test all of these expectations, we can do a basic analysis of tri-nucleotide sites within our sample of segregating sites.

As expected, the number of tri-nucleotide sites is higher for CpG sites (18,168 at CpG sites and 11,264 at non-CpG sites), as is the overall proportion of these sites within their respective type classes (0.7% at CpG sites and 0.2% at non-CpG sites; *χ*^2^ = 12,998, *p* < 2.2×10^−16^). Generally, the average segregating GC frequency at tri-nucleotide sites is high compared to di-nucleotide sites (Supplementary table 1). Furthermore, tri-nucleotides segregating at CpG sites tend to have higher GC frequency, compared to non-CpG sites (*t* = 25.089, *p* < 2.2×10^−16^). These observations indicate that sites segregating at high GC frequency are especially susceptible to additional mutations and formation of tri-nucleotide sites. Therefore, we expect a negative correlation between *B* values and tri-nucleotide abundance. Indeed, we observe a highly significant negative correlation between transversion *B* values and the number of tri-nucleotides (Spearman’s *ρ* = −0.8235, *p* = 0.0001), while the same is not observed for transitions (Spearman’s *ρ* = 0.2206, *p* = 0.4103; Figure 6). This suggests that, at least for rare CpG mutation types such as CpG transversions, the GC frequency distribution is affected by the general hypermutablity of CpG sites. Importantly, this observation does not exclude the action of other aforementioned modifiers of GC frequency distributions, which may interact to produce similar GC frequency patterns for other mutation types.

**Figure 6:**
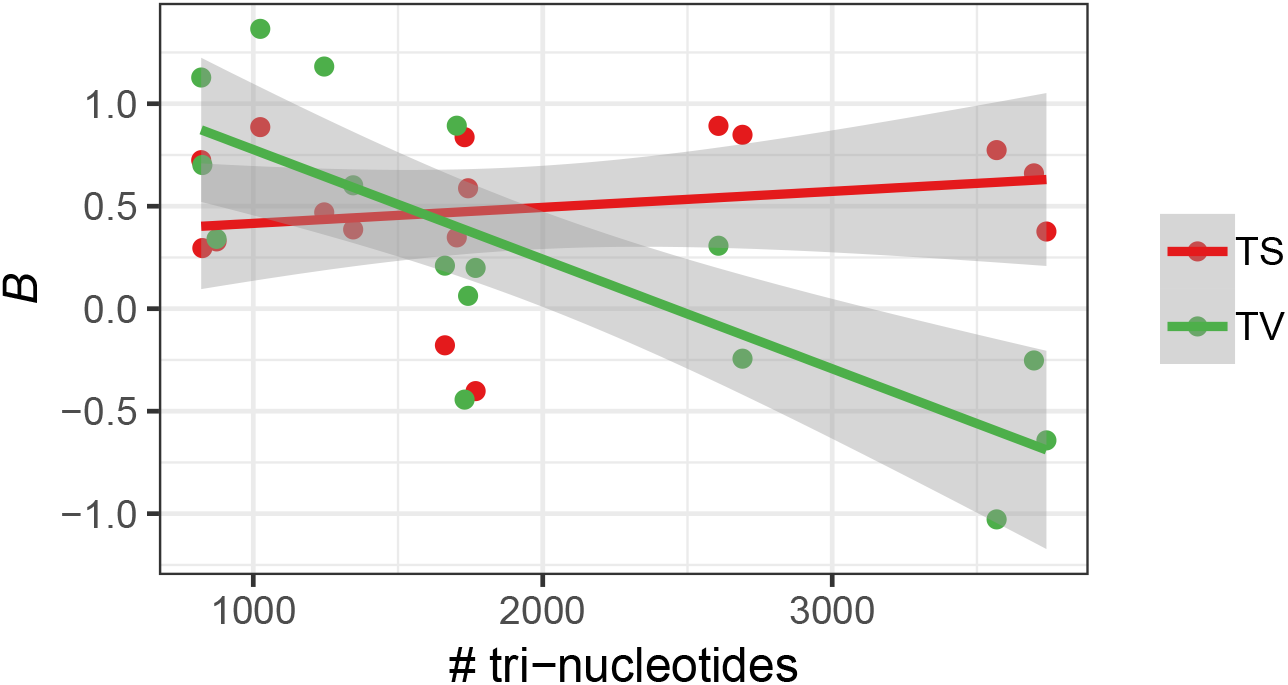
The correlation between the numbers of tri-nucleotide sites and *B* values for transitions and transversions of the YRI population. Each point represents a value for a specific flanking nucleotide context.

### 2.5 Fixation bias and recombination rate in great apes

In this section we investigate the GC fixation bias in eight great ape populations consisting of five subspecies of the *Pan* genus, one subspecies of the *Gorilla* genus and two subspecies of the *Pongo* genus. Additionally, we have derived recombination maps specific to each of these populations and study the relationship between the population-specific recombination rate and the fixation parameter *B*.

The distributions of per chromosome *B* estimates for the populations of the *Pan* genus are similar to those of humans, with CpG transitions having the highest values, followed by non-CpG transversions and transitions (Figure 7A). A direct comparison between per chromosome *B* values of the *P. paniscus* and human YRI population reveals no difference for CpG transitions (*t* = 0.0063, *p* = 0.995), reflective of similar effective sizes of the two populations [42] and a historically stable recombination environment experienced by this mutation type across the two species. On the other hand, somewhat higher *B* values are estimated for non-CpG transitions (*t* = 4.7911, *p* = 2.335×10^−5^) and transversions (*t* = 7.2361, *p* = 1.795×10^−8^) in humans, which may occur if these mutation types are affected by the differential evolution of the recombination landscape since their split from the common ancestor [36]. Notably, at the 1 Mb scale, the correlations between recombination rate and average GC frequency of segregating mutations for the *P. paniscus* population are positive (Spearman’s *ρ* = 0.3208, *p* < 2.2×10^−16^ for non-CpG transitions; Spearman’s *ρ* = 0.3614, *p* < 2.2×10^−16^ for CpG transitions; Spearman’s *ρ* = 0.2520, *p* < 2.2×10^−16^ for transversions; Supplementary figure 9A) and similar to the observed trend in humans (Figure 1B). Additionally, as in humans (Figure 1C), the correlations between recombination rate and mutation abundance are positive and strongest for CpG transitions (Spearman’s *ρ* = = 0.1739, *p* < 2.2×10^−16^ for non-CpG transitions; Spearman’s *ρ* = 0.5658, *p* < 2.2×10^−16^ for CpG transitions; Spearman’s *ρ* = 0.1568, *p* < 2.2×10^−16^ for transversions; Supplementary figure 9B). These results further demonstrate that large-scale gBGC dynamics are mostly congruent between chimpanzees and humans.

**Figure 7:**
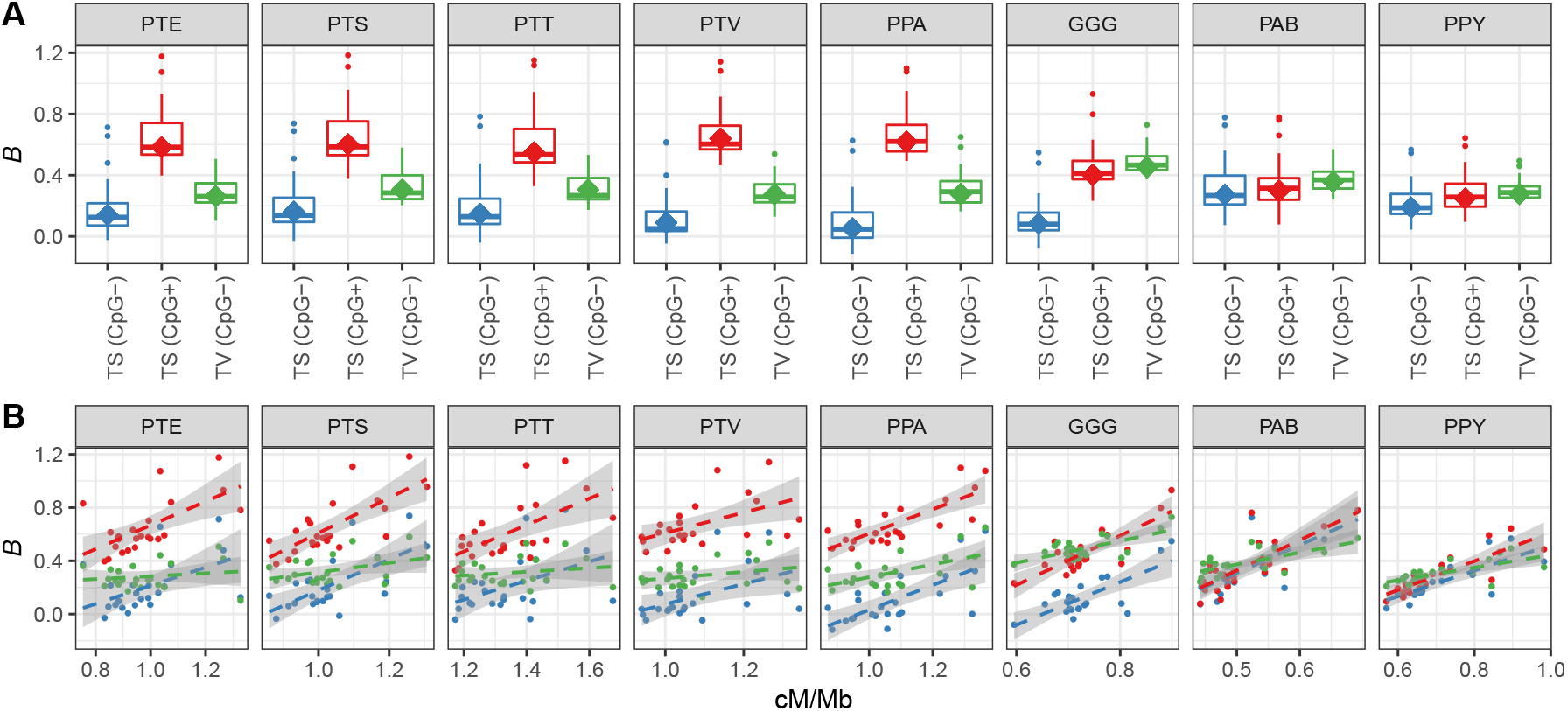
**A.** Distribution of the per-chromosome *B* parameter for non-CpG transitions (TS; CpG−), CpG transitions (TS; CpG+) and GC-changing transversions (TV; CpG−). The “diamond” points correspond to the genome-wide *B* estimated for all autosomes together. **B.** Relationship between the non-CpG transitions (blue), CpG transitions (red) or GC-changing transversions (green) per-chromsome *B* parameter and the sex-averaged recombination rate. The eight panels in each figure correspond to the *P. troglodytes ellioti* (PTE), *P. troglodytes schweinfurthii* (PTS), *P. troglodytes troglodytes* (PTT), *P. troglodytes verus* (PTV), *P. paniscus* (PPA), *G. gorilla gorilla* (GGG), *P. abelii* (PAB) and *P. pygmaues* (PPY) populations.

In contrast to humans and *Pan* species, great apes of the *Gorilla* and *Pongo* genus have the highest *B* estimates for non-CpG transversions, followed by CpG and non-CpG transitions. Therefore, the most notable characteristic of these species is the reduction of gBGC intensity at CpG transitions. Accordingly, we observe a reduced mean GC frequency of segregating CpG transitions (55.66% GC for *G. gorilla gorilla* and 53.21% GC for *P. abelii*) compared to non-CpG transversions (59.4% GC for *G. gorilla gorilla* and 61.4% GC for *P. abelii*) that is highly significant (*t* = 82.0, *p* < 2.2×10^−16^ for *G. gorilla gorilla*; *t* = 221.63, *p* < 2.2×10^−16^ for the *P. abelii*). At the 1 Mb scale, the correlations between recombination rate and average segregating GC frequency or mutation abundance for *Gorilla* and *Pongo* populations are positive (Supplementary figure 9C-F), but also notably different compared to the same correlations for human and *Pan* populations. Therefore, gBGC dynamics have likely diverged between humans and more distantly related great ape species. An important contributing factor to this observation could be differences in recombination landscapes between the species, as *Gorilla* and *Pongo* populations have more uniform recombination landscapes compared to other great apes [53] (Supplementary figure 2B). Such recombination landscapes may hypothetically result in more similar distributions of *B* values across different mutation categories for these populations (Figure 7A). Furthermore, great apes of the *Gorilla* and *Pongo* genus have the highest per chromosome *B* estimates for non-CpG transversions, while subspecies of the *Pongo* genus have the highest *B* values for non-CpG transitions, in accordance with the historically high effective sizes of these populations [42].

Similarly to human populations, the linear regression coefficients for the relationship between per chromosome *B* estimates and recombination rates are positive for all populations and mutation classes (Table 4). At the chormosome level, recombination rates explain ~ 16-60%, ~ 23-69% and ~ 25-48% of variance in *B* estimates for non-CpG transitions, CpG transitions and non-CpG transversions, respectively. These estimates overlap with those of human populations. Notably, the increase in *B* with unit increase in recombination rate is highest for the *Gorilla* and *Pongo* populations, which is likely due to their more uniform recombination rates (Supplementary figure 2B). When considering GC-changing mutations classified according to their flanking nucleotide composition, we observe similar patterns as in humans, especially for the *P. paniscus* population. The proportions of different mutation classes are similar between representative populations of each genus, with a general excess of CpG transitions and the highest paucity of CpG transversions (Figure 8A). Furthermore, between-species differences in *B* estimates across the 32 mutation categories are subtle, and major characteristics, such as the negative *B* values for CpG transversions are present in all species (Figure 8B). Notably, the majority of observed patterns in humans are also replicated in great apes, especially in the closest human relatives - bonobos and chimpanzees. Therefore, the differential impact of effective population sizes, recombination and gBGC on distinct mutation types is a conserved phenomenon of the *Hominidae* family.

**Figure 8:**
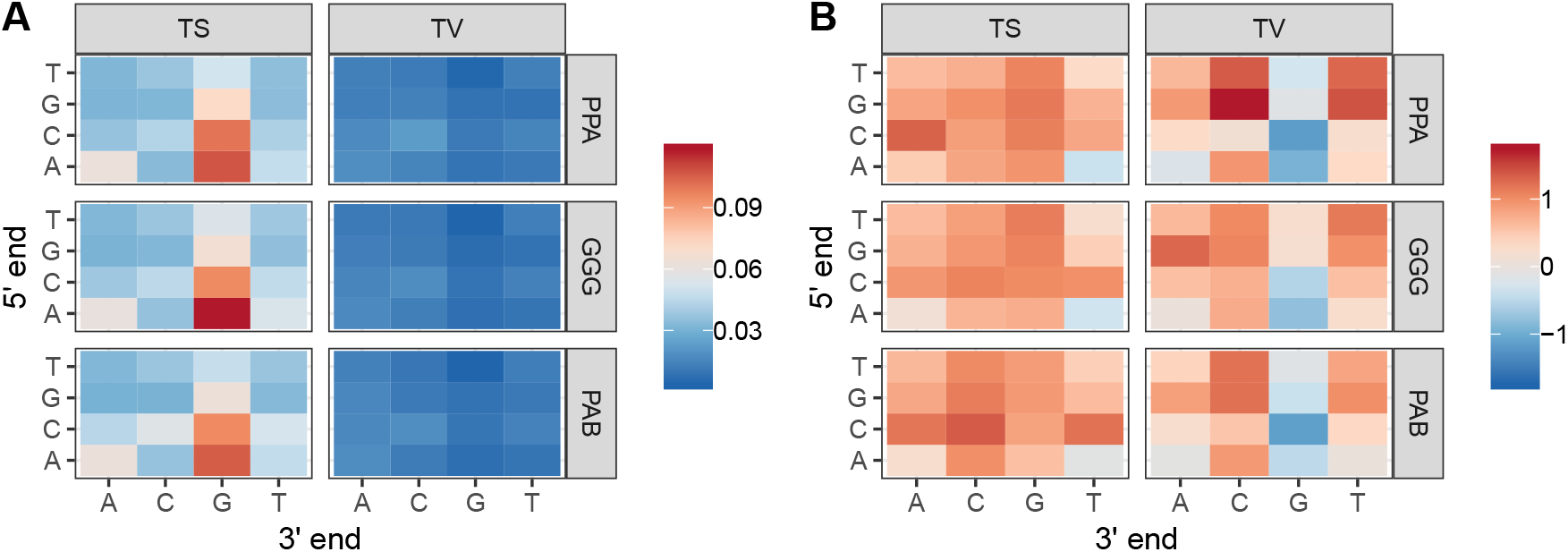
**A.** The proportions of the 32 different mutation types for the *P. paniscus* (PPA), *G. gorilla gorilla* (GGG) and *P. abelii* (PAB) populations. **B.** The values of the fixation bias *B* of 32 different mutation types for the PPA, GGG and PAB populations.

**Table 4:** Linear regression coefficients with corresponding *R*^2^ values for the relationship between per-chromsome *B* of non-CpG transitions (TS; CpG−), CpG transitions (TS; CpG+) or GC-changing transversions (TV; CpG−) and recombination rate, for different great ape populations (PTE = *P. troglodytes ellioti*; PTS = *P. troglodytes schweinfurthii*; PTT = *P. troglodytes troglodytes*; PTV = *P. troglodytes verus*; PPA = *P. paniscus*; GGG = *G. gorilla gorilla*; PAB = *P. abelii*; PPY = *P. pygmaues*).

| Parameter | Population |  |  |  |
| --- | --- | --- | --- | --- |
|  | PTE | PTS | PTT | PTV |
| $B(\text{TS; CpG-})$ | 0.6849 ($R^2 = 0.2134$)* | 1.1613 ($R^2 = 0.4117$***) | 0.7148 ($R^2 = 0.1621$*) | 0.7641 ($R^2 = 0.1999$*) |
| $B(\text{TS; CpG+})$ | 0.8927 ($R^2 = 0.3640$)** | 1.3206 ($R^2 = 0.5150$***) | 0.9952 ($R^2 = 0.3369$)** | 0.7729 ($R^2 = 0.2332$*) |
| $B(\text{TV; CpG-})$ | 0.1088 (*n.s.*) | 0.3525 (*n.s.*) | 0.1406 (*n.s.*) | 0.2455 (*n.s.*) |
| Parameter | PPA | GGG | PAB | PPY |
| $B(\text{TS; CpG-})$ | 0.9297 ($R^2 = 0.4688$***) | 1.6106 ($R^2 = 0.5122$***) | 2.0949 ($R^2 = 0.5026$***) | 0.9643 ($R^2 = 0.6006$***) |
| $B(\text{TS; CpG+})$ | 0.9110 ($R^2 = 0.5911$***) | 1.8518 ($R^2 = 0.6195$***) | 2.2734 ($R^2 = 0.5771$***) | 1.0836 ($R^2 = 0.6897$***) |
| $B(\text{TV; CpG-})$ | 0.4762 ($R^2 = 0.2692$)** | 0.8227 ($R^2 = 0.4167$***) | 0.8979 ($R^2 = 0.3951$***) | 0.4756 ($R^2 = 0.4276$***) |

\*\*\* $p < 0.001$ ; \*\* $p < 0.01$ ; \* $p < 0.05$ ; *n.s.* = non-significant

## 3 Discussion

Within this article, we use correlation analysis and population genetic theory to quantify the difference in gBGC dynamics between different types of GC-changing mutations. The basis of our analysis is the estimation of GC frequencies at segregating sites using large population genomic datasets, allowing us to examine the relationship between segregating GC content, recombination, and the gBGC parameter *B*. As the role of mutation processes on skews of GC frequency should be minor and transient, we conclude that differences in GC frequency distributions of different mutation types are largely due to the complex relationship between the effective population size, the distribution of different mutation types across the genome and the features of the recombination map, such as its rate intensity, sex-specificity, temporal stability and shape.

The stronger correlations between the average GC frequency and recombination for transitions, compared to the same correlation for transversions, indicates that transition-segregating sites are more prone to GC-biased gene conversion – a result that is in accordance with early molecular studies of MMR [4, 6, 60, 61]. Generally, mismatch repair is an essential process for maintaining genomic integrity. The most common types of mutations during replication are misincorporations of A or T nucleotides in place of G or C. Therefore, post-replicative MMR machinery may have evolved to preferentially recognize these mismatches and restore them back to G:C pairs. Since mismatches in recombination heteroduplexes are resolved using much of the molecular machinery of post-replicative MMR, recombination-associated gBGC may simply be an indirect consequence of the coevolution between the prevalence of AT-biased mutations and GC-biased MMR during DNA replication. Correspondingly, since transitions are more common than transversions, a higher specificity of MMR for transition-associated mismatches and consequently, a higher GC bias may be expected. Recent studies of non-crossover (NCO) gene conversions in a Mexican-American population estimated the GC-biased conversion ratio to be 0.7: 0.3 for transitions and 0.64: 0.36 for transversions [62], while a similar study in the Icelandic population resulted in ratios of 0.7: 0.3 and 0.68: 0.32 for transitions and transversions, respectively [20]. Given that the rate of NCO conversions is equal for both mutation types [20], the conversion ratios measured in humans would result in a 1.1-1.4× difference between transition and transversion *B* estimates. While tentative, this difference is not statistically significant. Furthermore, [20] detected a greater GC bias for CpG SNPs, in accordance with our results, while no such bias was detected by [62]. Given that the difference in the increase of average GC frequency per 1 cM/Mb is only ~ 2% between the two mutation types (Table 1), the number of conversion events from pedigree studies is likely not sufficient to detect such subtle frequency shifts.

Our estimates of the *B* parameter represent gBGC dynamics given the long-term, average *N_e_* of the studied populations. Therefore, if the conversion bias changed between transitions and transversions in the recent past, this may affect the transition-transversion difference derived from human pedigree studies compared to our estimated difference. Additionally, since the pedigree studies focused only on non-crossover conversions, there is a substantial fraction of crossover conversions that remain unexplored, which may further impede the comparison between gBGC dynamics inferred from pedigree data and the population genomic approach presented herein. Another possible source of uncertainty in our *B* estimates could be the fact that we do not consider the probability flux that arises when considering allele frequency spectra of multiple mutation types that have different mutation rate intensities [7]. As a consequence of this flux, AFSs may become skewed even in the absence of biased fixation dynamics. However, it has been shown that in the case of great apes such flux is inadequate when explaining the skewness of the AFS [46]. The difference between transition and transversion *B* estimates could also arise if *N_e_* along the genome varies locally with the distribution of transitions or transversions. Specifically, transversion mutations are likely to be more deleterious in both coding [64] and regulatory regions [19] and therefore experience stronger purifying selection, which would in turn reduce *N_e_* around transversions, more so than transitions. However, the stronger deleterious effect of transversions in regulatory and coding regions likely has a small effect on our estimates due to the fact that these mutations are only a small fraction of all segregating sites that we analyze.

Our analyses show that the major factor in determining the patterns of gBGC dynamics is the historical effective size of the population. This is most evident when comparing per chromosome *B* estimates for CpG transitions and non-CpG transversions between human populations, with Africans having higher values compared to Eurasians (Figure 3A and 4, Supplementary figure 5). Additionally, great ape populations with high effective sizes of the *Gorilla* and *Pongo* genus have the highest *B* estimates for non-CpG transversions and non-CpG transitions, respectively (Figure 7A).

However, there are notable discrepancies in opposition to this pattern, that are likely due to factors other than effective size differences. Specifically, Europeans have higher *B* estimates for non-CpG transitions compared to both Africans and Asians, while *B* estimates of CpG transitions for *Gorilla* and *Pongo* populations are notably lower compared to other species. Such patterns are likely due to between-species differences in recombination landscapes (Supplementary Figure 2) and/or shifts in distributions and frequencies of different mutation types [23]. Additionally, the analysis of male and female recombination maps (Table 2) implies that sex-specific meiotic processes are important determinants of gBGC dynamics. This observation could be due to sex-specific differences of the meiotic process, involving differences in the duration of meiotic phases, crossover rates and positions, and possibly repair mechanisms involved in sex-specific mismatch resolution.

When considering gBGC dynamics of GC-changing mutations that have been classified into categories according to their flanking nucleotide context, we uncovered that some mutation categories have negative *B* values, in opposition to the gBGC hypothesis. This is a surprising result, that is likely due to CpG hypermutability in the case of transversions segregating at CpG sites (Figure 6). However, negative *B* values for 5’ACT3’↔5’ATT3’ and 5’CCT3’↔5’CAT3’ mutation types have likely arisen by different mechanisms, as they cannot be explained by CpG hypermutability. Therefore, more investigation is needed into the dependence of gBGC dynamics on flanking nucleotide context.

We illustrate the importance of analyzing population genomic datasets with their corresponding population-specific recombination maps when considering gBGC dynamics. Given the current accumulation of large population genomic datasets that are suitable for recombination map estimation, recombination-associated processes can now be studied at an unprecedented scale, providing us with a better understanding of the effect of fixation biases on the evolution of nucleotide composition across a large variety of species.

## 4 Conclusions

Our study shows that gBGC dynamics differ among mutation types. Transitions and transversions differ in their genomic distributions which alter the average gBGC intensity experienced by these mutations. Most notably, CpG transitions that are enriched in highly recombining, GC-rich regions are most strongly affected by gBGC. An important modifier of gBGC efficiency is the effective size of the population, but not all gBGC-related patterns are in accordance with population sizes, as demonstrated by non-CpG transition dynamics in European populations and CpG transition dynamics in *Gorilla* and *Pongo* populations. Additionally, we show that sex specificity of recombination maps and flanking nucleotide context play important roles in determining gBGC dynamics of different mutation types. Therefore, by explicitly considering different mutation types, we uncover novel aspects of nucleotide composition evolution that serve to deepen our knowledge of basic evolutionary processes.

## 5 Methods and Materials

### 5.1 Data

The single nucleotide polymorphism (SNP) data for the Yoruba (YRI), Toscana (TSI) and Han Chinese (CHB) populations were taken from the 1000 Genomes Project Consortium [10] and the corresponding recombination maps from [50]. The SNP data for the Icelandic population (ISL) were taken from [27] and the recombination maps from [21]. The great ape dataset was curated from four different sequencing studies [12, 38, 42, 63], consisting of samples from five subspecies of the *Pan* genus: *P. troglodytes ellioti* (PTE; *N* = 10), *P. troglodytes schweinfurthii* (PTS; *N* = 19), *P. troglodytes troglodytes* (PTT; *N* = 18), *P. troglodytes verus* (PTV; *N* = 11) and *P. paniscus* (PPA; *N* = 13); one subspecies of the *Gorilla* genus: *G. gorilla gorilla* (GGG; *N* = 23); and two subspecies of the *Pongo* genus: *P. abelii* (PAB; *N* = 11) and *P. pygmaeus* (PPY; *N* = 15).

### 5.2 Estimation of GC frequency and allele frequency spectra

We conducted all analyses after filtering out singleton sites and sites segregating at frequency < 0.5%. For all analyses, we excluded sites masked by RepeatMasker [49] in the reference sequence. For the regression analyses, we only considered non-overlapping autosomal windows with at least 100 segregating sites and an average recombination rate of > 0 cM/Mb and < 5 cM/Mb. Due to the large sample sizes, allele frequency spectra (AFSs) for the human populations were constructed by classifying sites into 50 equally-sized GC frequency bins.

To estimate GC frequencies of segregating sites for the curated great ape dataset, we conducted a *de novo* read mapping and variant calling procedure for all eight studied great ape populations. The reference genomes used for mapping short reads were Clint PTRv2 for chimpanzee (University of Washington; January 2018), gorGor4 for gorilla (Wellcome Trust Sanger Institute; October 2015) and Susie PABv2 for orangutan (University of Washington, 2018); https://www.ncbi.nlm.nih.gov/. The short read data were mapped using the bwa mapper, version 0.7.5a [29]. Filtering and merging of bam files was done using sambamba v0.5.1 [55]. We retained properly paired reads with mapping quality MQ ≥ 50 and less than 2 mismatches to the reference, and filtered out duplicated, unmapped and secondarily aligned reads. The variants were called separately for each individual sample using HaplotypeCaller in −ERC GVCF mode of gatk v4.0 [31]. Genotypes were called jointly for the respective subspecies populations using gatk GenotypeGVCFs. The called SNPs were filtered according to gatk guidelines for hard filtering (-filter “QD < 2.0 —— FS > 60.0 —— MQ < 40.0 —— MappingQualityRankSum < −12.5 —— ReadPosRankSum < −8.0”). Furthermore, vcftools v0.1.16 [11] was used to filter out sites where more than 20% of individuals had missing data and/or sites which failed the Hardy-Weinberg equilibrium test (*p* < 0.001). We then filtered out SNPs covered by less than 1.5× n reads (where n is the total ploidy of the region, thus ensuring that, on average, SNPs are covered by at least 1.5 reads per haploid genome) or more than 2×n×cov reads (where cov is the mean coverage per haploid genome). As with human datasets, we excluded sites masked by RepeatMasker [49] in the respective reference sequences. Due to the small samples sizes of great ape populations we assigned GC-changing sites into discrete GC frequency categories to construct the corresponding AFSs.

### 5.3 Estimation of the *B* parameter

Following [58], we consider a Moran model of evolution for a biallelic locus under the influence of mutation (*θ* = *N μ*) and fixation bias towards the preferred alleles (*B* = *N_e_b*). In our application, *B* quantifies the preferred fixation of GC over AT alleles. Under mutation-fixation-drift equilibrium, and assuming low mutation rates and a high sample size, the frequency *x* at which the preferred allele segregates in the population is

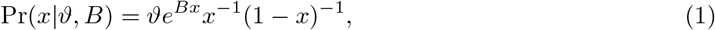

where *ϑ* = (*αβθ*) / (*αe^B^* + *β*) (*α* is a parameter ranging from 0 to 1 which quantifies the mutation bias towards the preferred allele; *β* =1 – *α* is the mutation bias towards the unpreferred allele). Assuming binomial sampling, the probability that a SNP is present at frequency *y* in a sample of *M* haploid sequences is

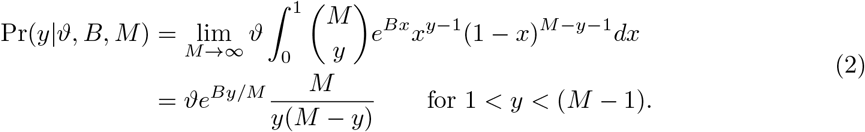

This formula is a good approximation if *αβθ* < 0.01 [59] (well within the range for mammals) and a sample of size *M* > 20 from a population evolving under equilibrium dynamics. Furthermore, we can treat *y* as a discrete frequency category or as a category encompassing a specific frequency range.

To account for the potential effect of demography and population structure on Pr(*y*|*ϑ, B, M*) we introduce nuisance parameters *r_y_* that are obtained by comparing the mutation-drift equilibrium expectation of the AFS with the empirical AFS of putatively neutral sites. In our case, we use the AFS from GC-conservative sites, *i.e*. sites segregating for A(T):T(A) or G(C):C(G) polymorphisms, as proxy for putatively neutral sites and define

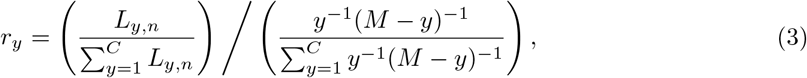

where *C* is the number of frequency categories of the AFS and *L_y,n_* is the count of GC-conservative sites in frequency category *y* of the neutral AFS. As these sites are unaffected by GC fixation bias, the choice of assigning them into the *y*th or (*M* – *y*)th frequency category of the AFS is random. We then normalize (2) with the sum across *y* categories and express the likelihood of the non-neutral AFS as

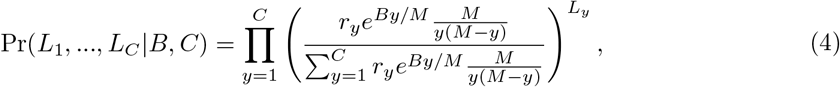

where *L_y_* is the count of GC-changing sites within the *y*th category of the non-neutral AFS. The estimate of *B* can be obtained by maximizing the likelihood (4) as described in Appendix A of [58]. Furthermore, we omit the lowest and highest categories of the AFS before estimating *B*, since singleton sites assigned to these extreme categories are likely to deviate most from equilibrium conditions and therefore bias our estimation method.

### 5.4 Classification of mutations according to flanking context

Each GC-changing mutation is classified into a mutation type category according to the 5’- and 3’-adjacent base. We exclude segregating sites that are adjacent to one another. Additionally, we group together mutation types that are reverse complements of each other. For example, a site segregating for a 5’ACG3’↔5’ATG3’ mutation is grouped together with site segregating for a 5’CGT3’↔5’CAT3’ mutation. This procedure results in 32 categories of GC-changing mutations, 16 categories each for transitions and GC-changing transversions.

### 5.5 Estimation of great ape recombination maps

Steps involved in recombination rate inference were adopted from previous studies that used similar data for map estimation [2, 53]. Briefly, to improve genotype calls, we first imputed the missing data using fastPHASE v1.4 [45] as described in [53] and then re-phased the inferred haplotypes with PHASE v2.1 [52] as described in [2]. Recombination maps were generated using the LDhat v2.2 program in interval mode [33]. Specifically, haplotypes were split into windows containing 4000 non-singleton SNPs with a 200 SNP overlap between windows. Recombination maps were inferred for each window separately and subsequently joined into chromosome-level recombination maps as described in [2]. To avoid potential biases in map estimation, we excluded data of four individuals from the *G. gorilla gorilla* subspecies population (Bulera, Kowali, Suzie and Oko) due to their high level of relatedness with other members of this population [42]. We converted the 4*N_e_r*/bp estimates of recombination rate provided by the LDhat v2.2 program into cM/Mb for each 1 Mb region by dividing the average 4*N_e_r*/bp of the region with the per base pair estimate of Watterson’s *θ_w_* across the 1 Mb region, normalized by the number of callable sites within the region. A site was defined as callable if covered by at least 1.5 reads per haploid genome and no more than 2×n×cov reads. Since *E*[*θ_w_*] = 4*N_e_μ* (where *μ* is the per base and generation mutation rate), the division yields (4*N_e_r*/bp)/*θ_w_* = *r*/*μ*. We then used the recently obtained estimate of *μ* = 1.25 × 10^−8^ for great apes [3] to obtain the probability of recombination r and obtain average estimates of recombination rates in 1 Mb regions as cM/Mb = 100*r*/10^6^.

## 6 Acknowledgments

We thank Marjolaine Rousselle and Claus Vogl for critical reading of the manuscript. The study was supported by grant NNF18OC0031004 from the Novo Nordisk Foundation.

## 7 Supplement

**Supplementary table 1:** Count and mean/median GC frequency of bi-allelic transitions and transversion, and tri-nucleotides segregating at non-CpG and CpG sites in the African YRI population.

|  | #CpG− | #CpG+ | mean/median GC (CpG−) | mean/median GC (CpG+) |
| --- | --- | --- | --- | --- |
| TS | 4,231,805 | 2,212,779 | 54.90%/67.59% | 60.01%/79.17% |
| TV | 1,269,834 | 250,334 | 58.21%/75.93% | 40.27%/21.76% |
| Tri-nucleotides | 11,264 | 18,168 | 62.16%/80.56% | 72.96%/89.81% |

**Supplementary figure 1:**
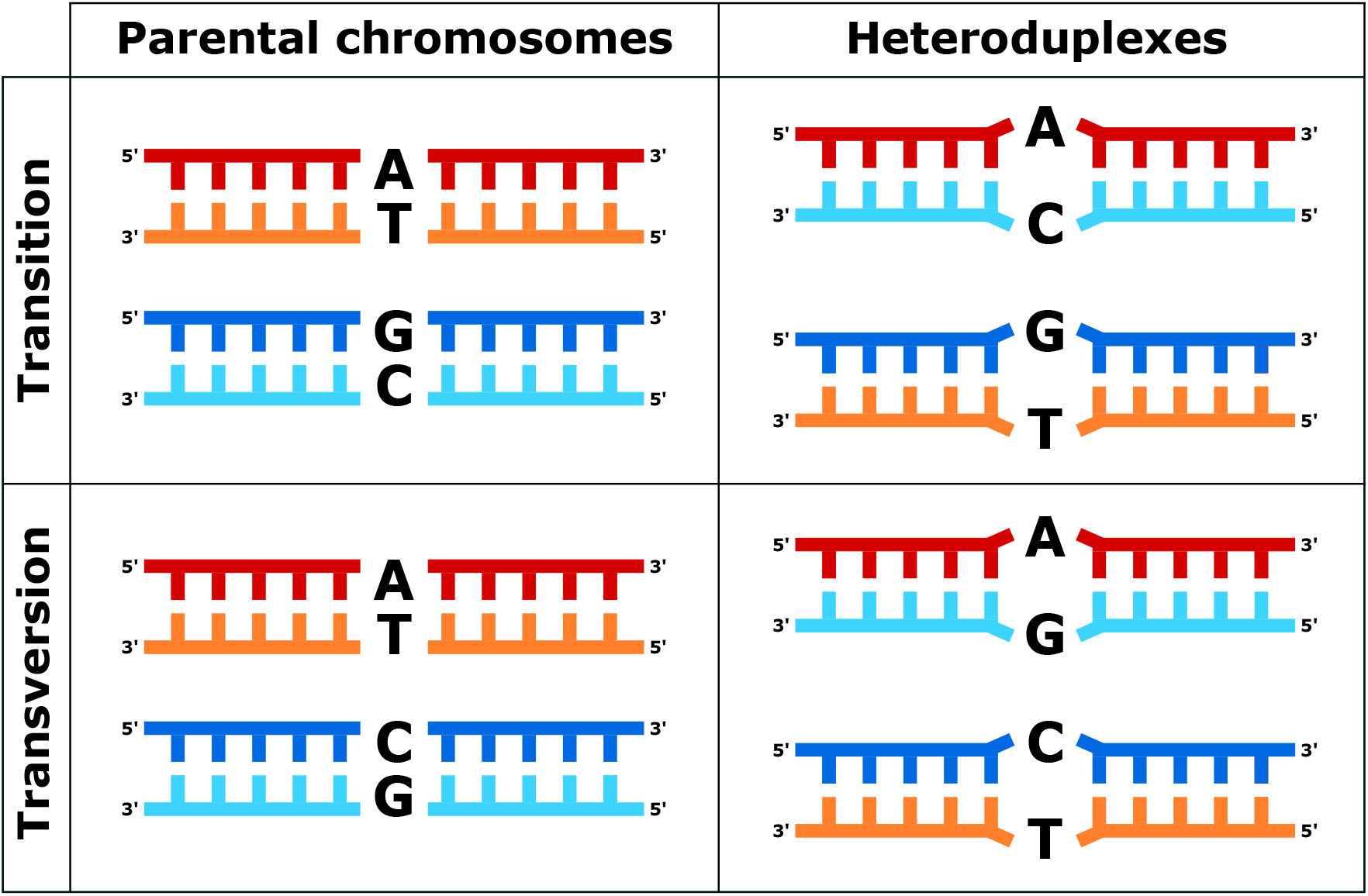
Two types of GC-changing mutations and the corresponding heteroduplex mismatches formed during recombination.

**Supplementary figure 2:**
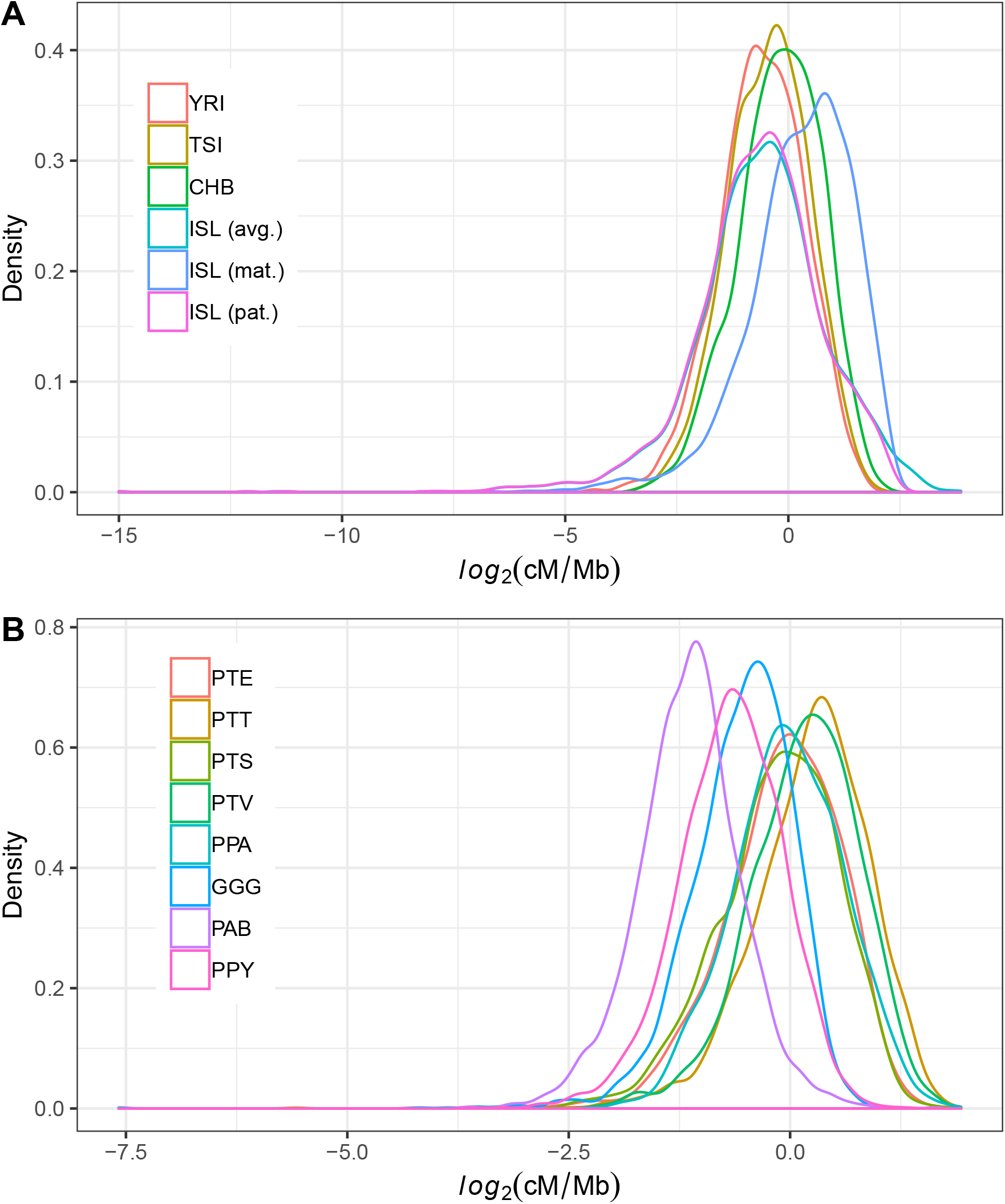
**A.** Distributions of average recombination rates in 1 Mb genomic regions for the Yoruba (YRI), Toscana (TSI), Han Chinese (CHB) and Iceland (ISL) populations. For the ISL, three different maps are available: sex-averaged (avg.), maternal (mat.) and paternal (pat.) map. **B.** Distributions of average recombination rates in 1 Mb genomic regions for different great ape populations (PTE = *P. troglodytes ellioti*; PTS = *P. troglodytes schweinfurthii*; PTT = *P. troglodytes troglodytes*; PTV = *P. troglodytes verus*; PPA = *P. paniscus*; GGG = *G. gorilla gorilla*; PAB = *P. abelii*; PPY = *P. pygmaues*).

**Supplementary figure 3:**
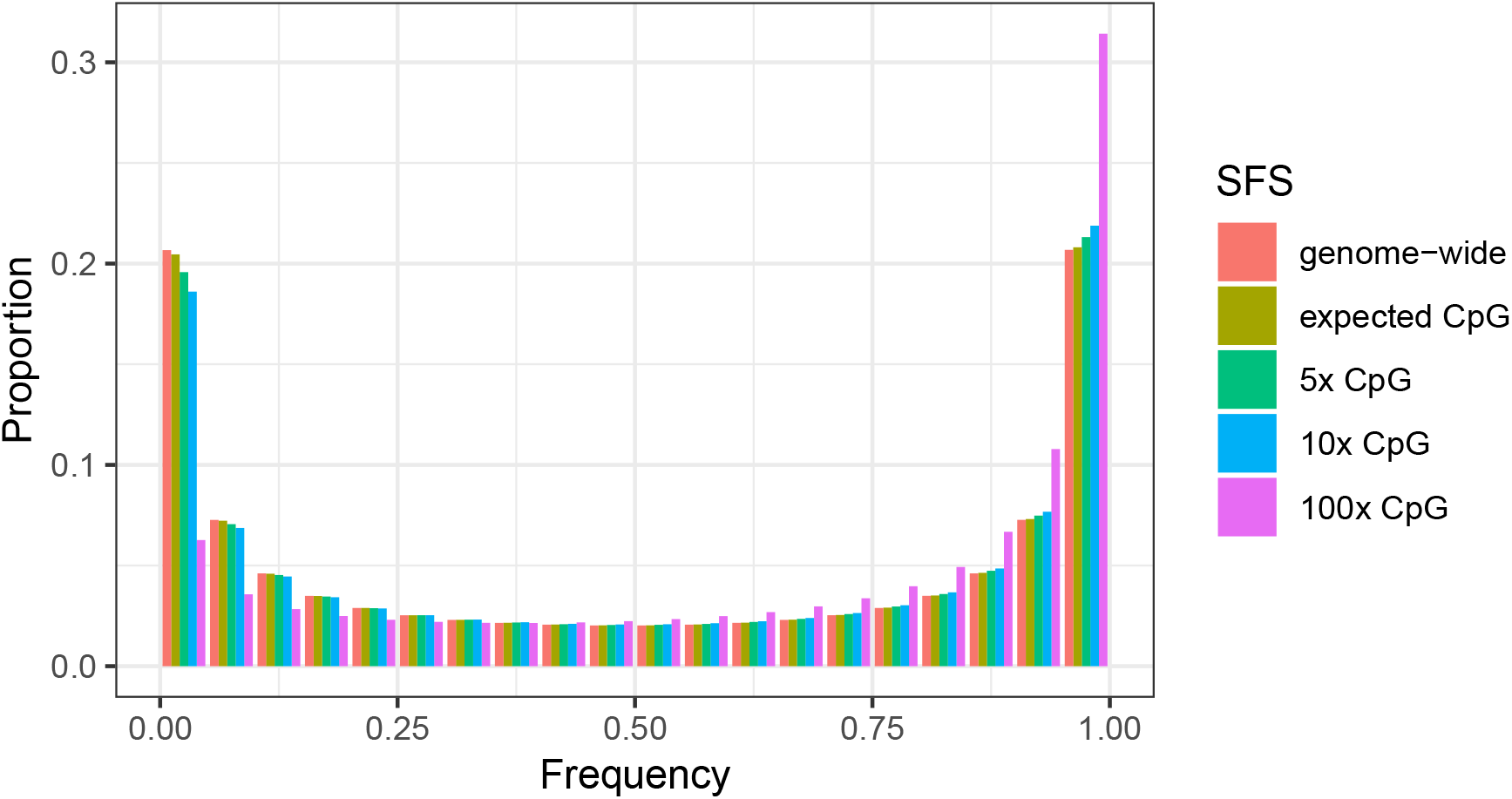
Expected SFS given a mutation-drift model and different rates of mutation, including the genome-wide mutation rate, CpG-specific mutation rate and multipliers of the CpG-specific mutation rate.

**Supplementary figure 4:**
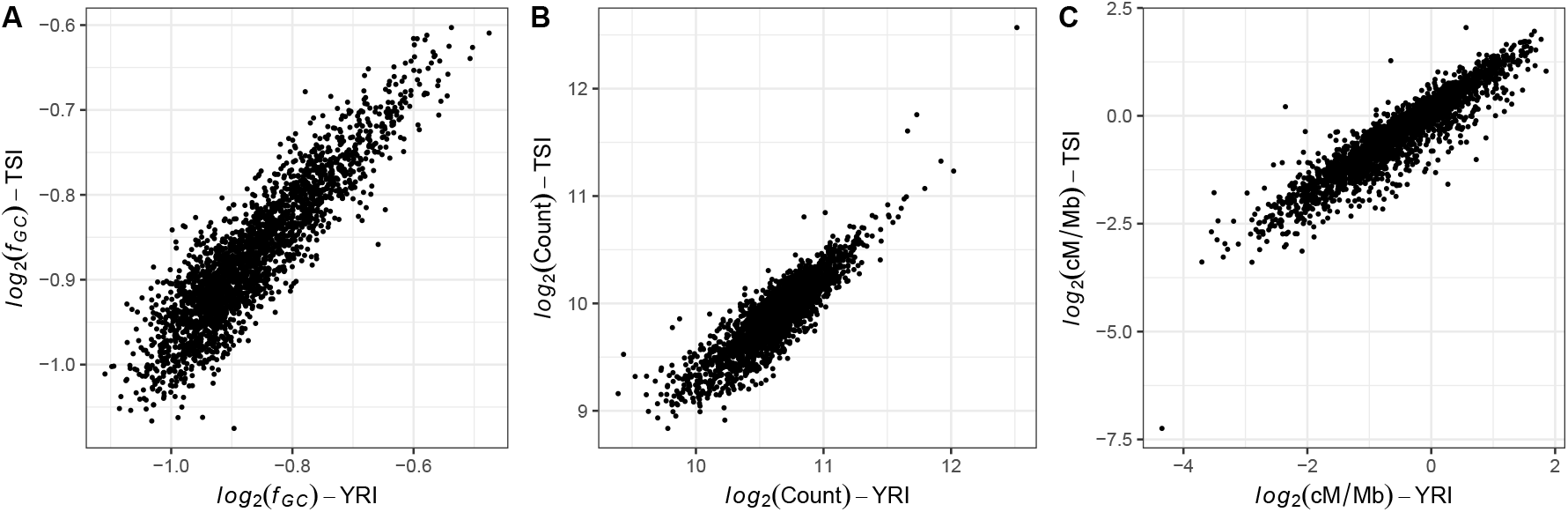
**A.** Correlation between the average GC frequency (*f*_GC_) of non-CpG transitions in 1 Mb autosomal windows of the African YRI and European TSI populations. **B.** Correlation between the count of non-CpG transitions in 1 Mb autosomal windows of the African YRI and European TSI populations. **C.** Correlation between the average recombination rate in 1 Mb autosomal windows of the African YRI and European TSI populations.

**Supplementary figure 5:**
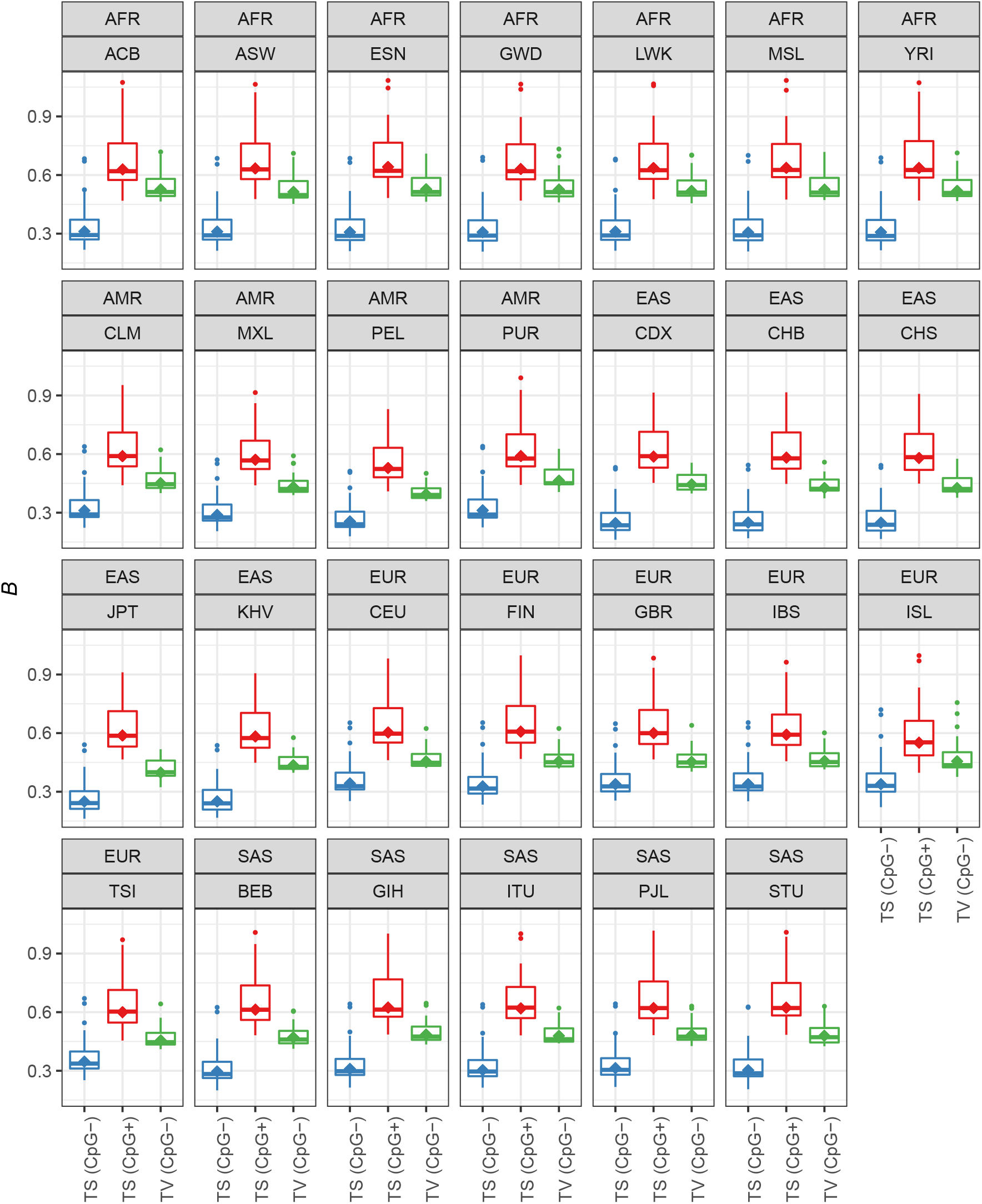
Distribution of the *B* parameter for non-CpG transitions (TS; CpG−), CpG transitions (TS; CpG+) and transversions (TV; CpG−) across autosomes for 26 human populations of the 1000 Genomes dataset and the Iceland population (ISL). In addition to specific population identifiers, we also indicate geographic superpopulation identifiers (AFR = African; SAS = South Asian; EAS = East Asian; EUR = European; AMR = Admixed American).

**Supplementary figure 6:**
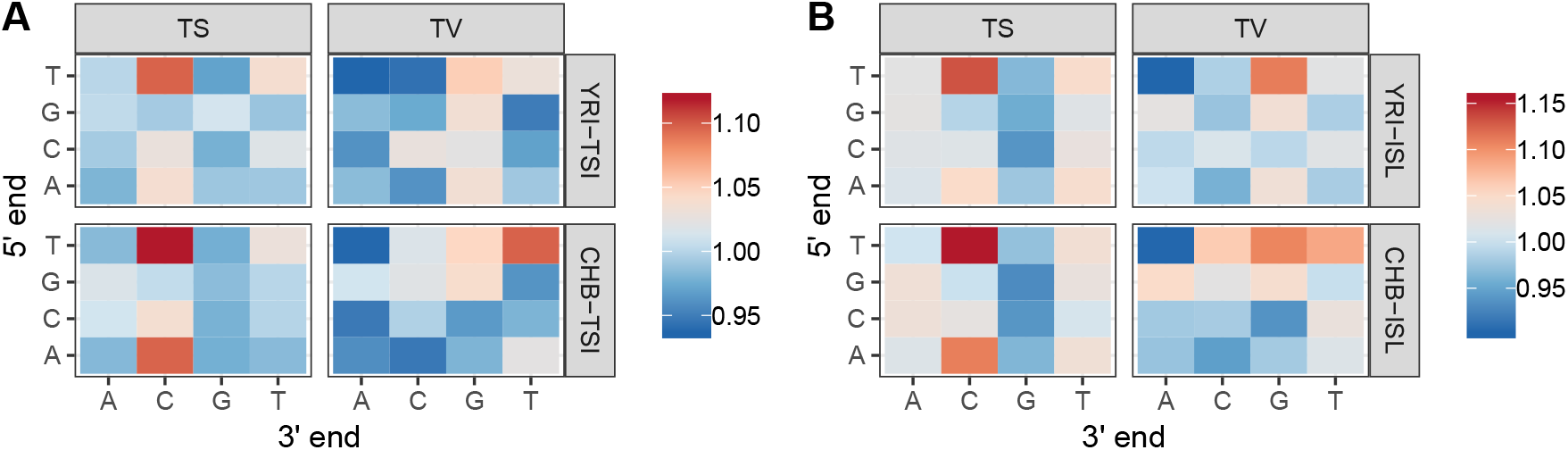
The ratios of proportions of the 32 different mutation types between the non-European populations and European **A.** TSI and **B.** ISL populations.

**Supplementary figure 7:**
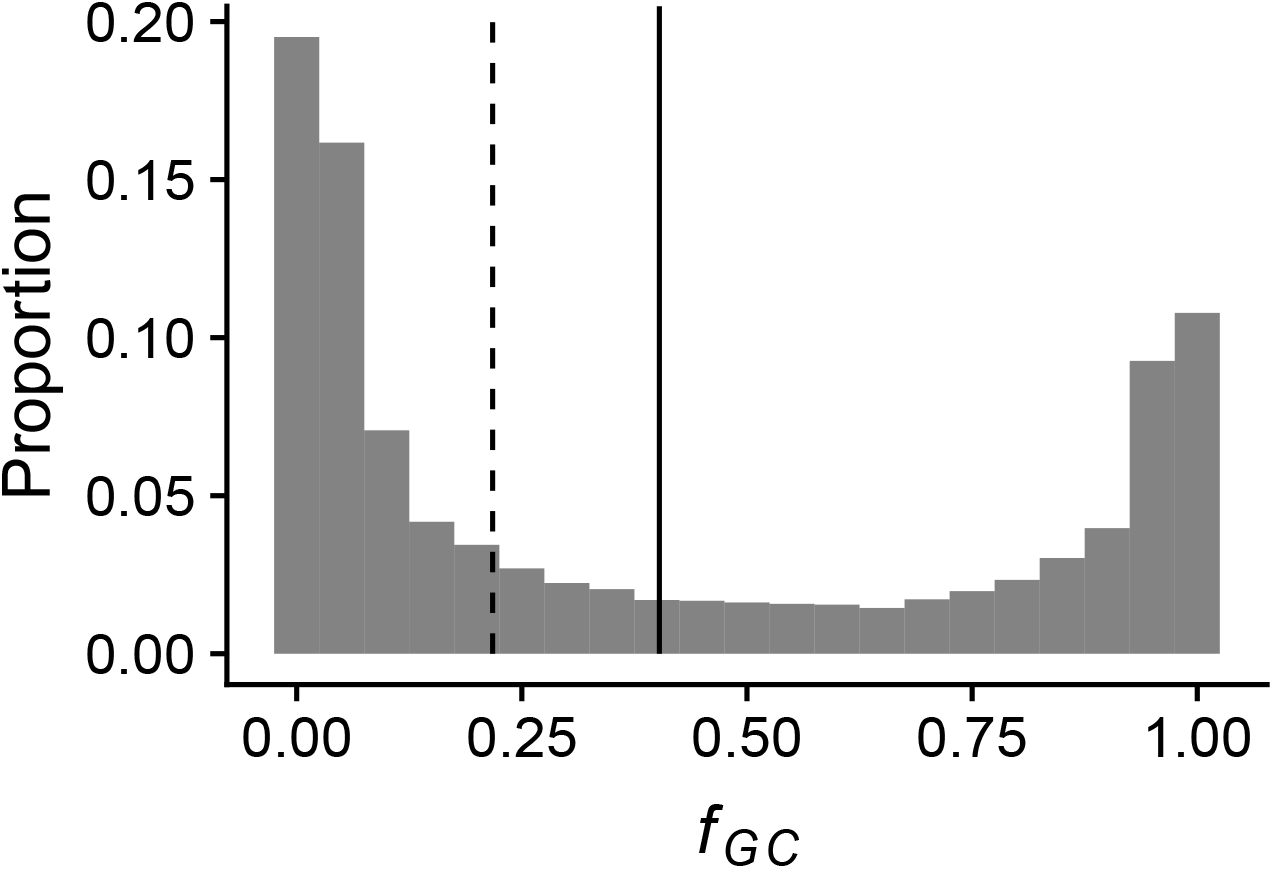
The distribution of GC frequencies (*f*_GC_) for CpG transversions of the YRI populations.

**Supplementary figure 8:**
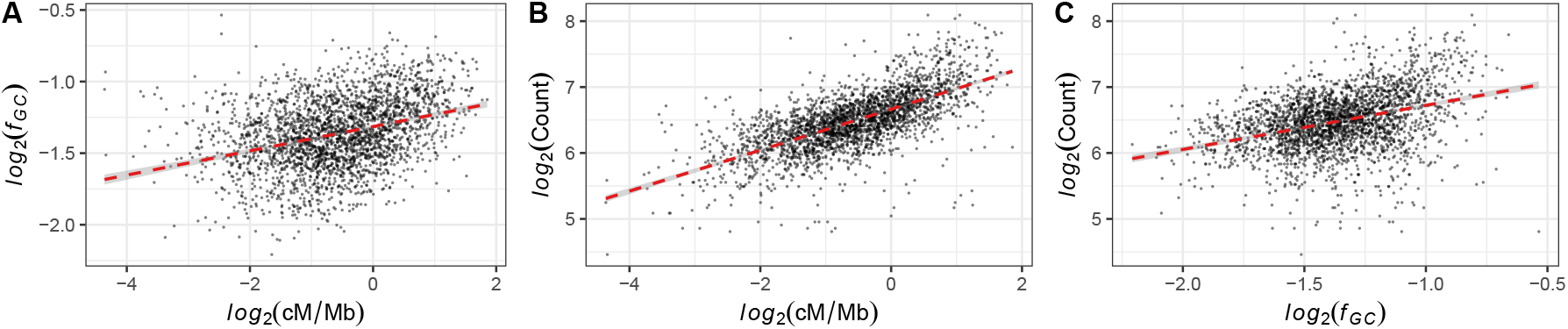
**A.** Relationship between the average GC frequency (*f*_GC_) at segregating CpG transversion sites and average recombination rate in 1 Mb autosomal windows of the African Yoruba population. **B.** Relationship between the count of CpG transversions and recombination rate in 1 Mb autosomal windows of the African Yoruba population. **C.** Relationship between the count of CpG transversion and average GC frequency (*f*_GC_) in 1 Mb autosomal windows of the African Yoruba population.

**Supplementary figure 9:**
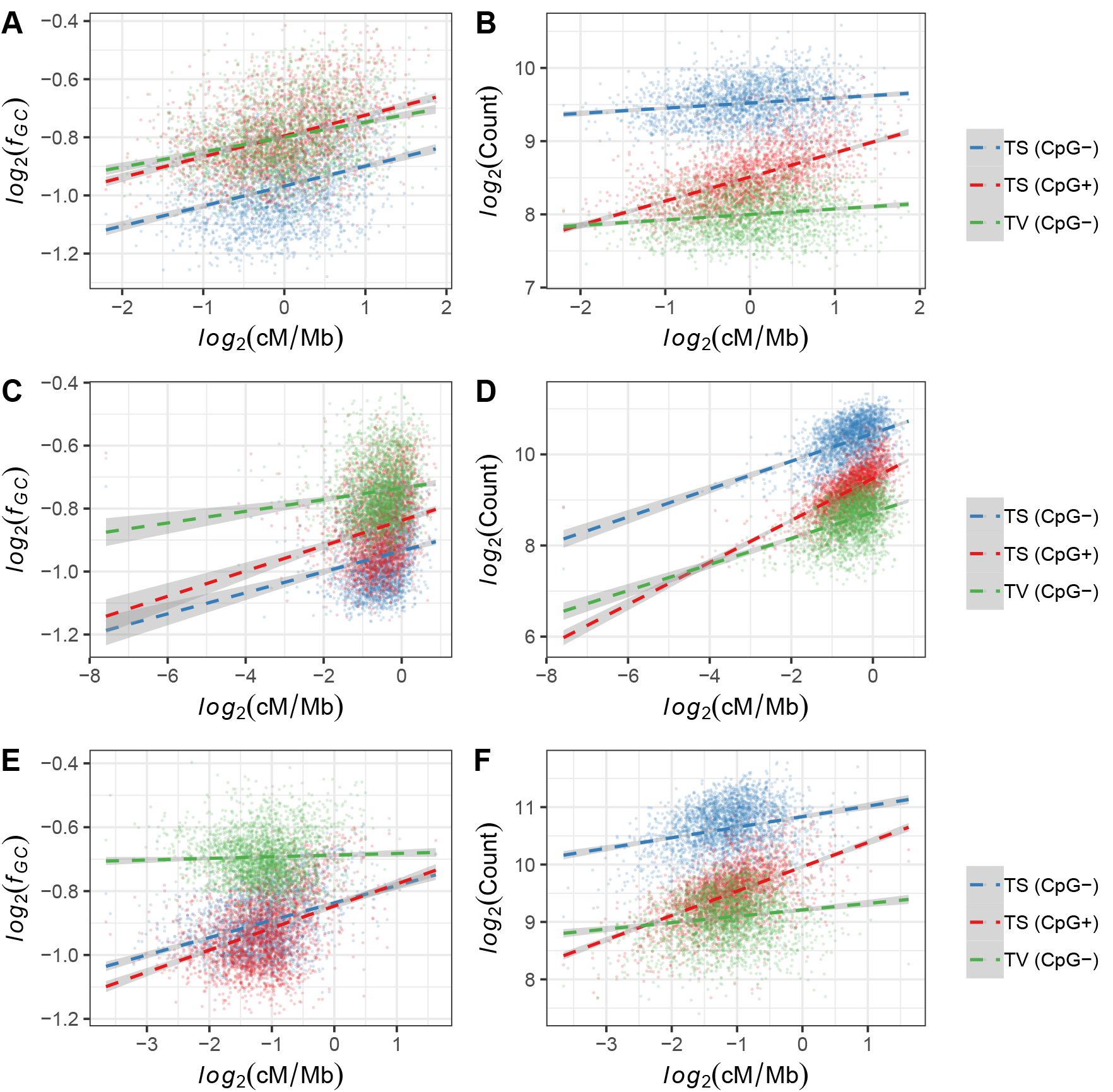
Relationship between the average GC frequency (*f*_GC_) at segregating sites and average recombination rate in 1 Mb autosomal for non-CpG transitions (TS; CpG−), CpG transitions (TS; CpG+) and GC-changing transversions (TV; CpG−) of **A.** the *P. paniscus* population, **C.** the *G. gorilla gorilla* population, and **E.** the *P. abelii* population. Relationship between the count of GC-changing segregating sites and average recombination rate in 1 Mb autosomal for non-CpG transitions (TS; CpG−), CpG transitions (TS; CpG+) and GC-changing transversions (TV; CpG−) of **B.** the *P. paniscus* population, **D.** the *G. gorilla gorilla* population, and **F.** the *P. abelii* population.

## References

[1] Barbara Arbeithuber, Andrea J. Betancourt, Thomas Ebner, and Irene Tiemann-Boege. Crossovers are associated with mutation and biased gene conversion at recombination hotspots. Proceedings of the National Academy of Sciences, 112(7):2109–2114, 2015.

[2] Adam Auton, Adi Fledel-Alon, Susanne Pfeifer, Oliver Venn, Laure Ségurel, Teresa Street, Ellen M. Leffler, Rory Bowden, Ivy Aneas, John Broxholme, Peter Humburg, Zamin Iqbal, Gerton Lunter, Julian Maller, Ryan D. Hernandez, Cord Melton, Aarti Venkat, Marcelo A. Nobrega, Ronald Bontrop, Simon Myers, Peter Donnelly, Molly Przeworski, and Gil McVean. A fine-scale chimpanzee genetic map from population sequencing. Science, 336(6078):193–198, 2012.

[3] Søren Besenbacher, Christina Hvilsom, Tomas Marques-Bonet, Thomas Mailund, and Mikkel Heide Schierup. Direct estimation of mutations in great apes reconciles phylogenetic dating. Nature Ecology & Evolution, 3(2):286–292, 2019.

[4] Colin A. Bill, Walter A. Duran, Nathan R. Miselis, and Jac A. Nickoloff. Efficient repair of all types of single-base mismatches in recombination intermediates in chinese hamster ovary cells: competition between long-patch and gt glycosylase-mediated repair of gt mismatches. Genetics, 149(4):1935–1943, 1998.

[5] Rui Borges, Gergely Szöllősi, and Carolin Kosiol. Quantifying gc-biased gene conversion in great ape genomes using polymorphism-aware models. Genetics, pages genetics–302074, 2019.

[6] Thomas C. Brown and Josef Jiricny. Different base/base mispairs are corrected with different efficiencies and specificities in monkey kidney cells. Cell, 54(5):705–711, 1988.

[7] Conrad J. Burden and Yurong Tang. Rate matrix estimation from site frequency data. Theoretical Population Biology, 113:23–33, 2017.

[8] Jedidiah Carlson, Adam E. Locke, Matthew Flickinger, Matthew Zawistowski, Shawn Levy, Richard M. Myers, Michael Boehnke, Hyun Min Kang, Laura J. Scott, Jun Z. Li, Sebastian Zöllner, Devinand Absher, Huda Akil, Gerome Breen, Margit Burmeister, Sarah Cohen-Woods, William G. Iacono, James A. Knowles, Lisa Legrand, Qing Lu, Matthew McGue, Melvin G. McInnis, Carlos N. Pato, Michele T. Pato, Margarita Rivera, Janet L. Sobell, John B. Vincent, Stanley J. Watson, and The BRIDGES Consortium. Extremely rare variants reveal patterns of germline mutation rate heterogeneity in humans. Nature Communications, 9(1):1–13, 2018. URL https://www.nature.com/articles/s41467-018-05936-5.

[9] Ujani Chakraborty and Eric Alani. Understanding how mismatch repair proteins participate in the repair/anti-recombination decision. FEMS Yeast Research, 16(6):fow071, 2016. URL https://academic.oup.com/femsyr/article/16/6/fow071/2198065.

[10] The 1000 Genomes Project Consortium. A global reference for human genetic variation. Nature, 526(7571):68–74, 2015.

[11] Petr Danecek, Adam Auton, Goncalo Abecasis, Cornelis A. Albers, Eric Banks, Mark A. DePristo, Robert E. Handsaker, Gerton Lunter, Gabor T. Marth, Stephen T. Sherry, Gilean McVean, Richard Durbin, and 1000 Genomes Project Analysis Group. The variant call format and vcftools. Bioinformatics, 27(15):2156–2158, 2011. URL https://academic.oup.com/bioinformatics/article/27/15/2156/402296.

[12] Marc de Manuel, Martin Kuhlwilm, Peter Frandsen, Vitor C Sousa, Tariq Desai, Javier Prado-Martinez, Jessica Hernandez-Rodriguez, Isabelle Dupanloup, Oscar Lao, Pille Hallast, Joshua M. Schmidt, José María Heredia-Genestarl, Andrea Benazzo, Guido Barbujani, Benjamin M. Peter, Lukas F. K. Kuderna, Ferran Casals, Samuel Angedakin, Mimi Arandjelovic, Christophe Boesch, Hjalmar Kühl, Linda Vigilant, Kevin Langergraber, John Novembre, Marta Gut, Ivo Gut, Arcadi Navarro, Frands Carlsen, Aida M. Andrés, Hans. R. Siegismund, Aylwyn Scally, Laurent Excoffier, Chris Tyler-Smith, Sergi Castellano, Yali Xue, Christina Hvilsom, and Tomas Marques-Bonet. Chimpanzee genomic diversity reveals ancient admixture with bonobos. Science, 354(6311):477–481, 2016.

[13] Christiane Dohet, Robert Wagner, and Miroslav Radman. Repair of defined single base-pair mismatches in escherichia coli. Proceedings of the National Academy of Sciences, 82(2):503–505, 1985. URL https://www.pnas.org/content/82/2/503.

[14] Adam Eyre-Walker. Recombination and mammalian genome evolution. Proceedings of the Royal Society of London. Series B: Biological sciences, 252(1335):237–243, 1993. URL https://royalsocietypublishing.org/doi/10.1098/rspb.1993.0071.

[15] Stephanie M. Fullerton, Antonio Bernardo Carvalho, and Andrew G. Clark. Local rates of recombination are positively correlated with gc content in the human genome. Molecular Biology and Evolution, 18(6):1139–1142, 2001. URL https://academic.oup.com/mbe/article/18/6/1139/1046938.

[16] Nicolas Galtier. Gene conversion drives gc content evolution in mammalian histones. Trends in Genetics, 19(2):65–68, 2003. URL https://academic.oup.com/mbe/article/21/7/1438/1080510.

[17] Nicolas Galtier, Gwenael Piganeau, Dominique Mouchiroud, and Laurent Duret. Gc-content evolution in mammalian genomes: the biased gene conversion hypothesis. Genetics, 159(2): 907–911, 2001. URL https://www.genetics.org/content/159/2/907.

[18] Sylvain Glémin, Peter F. Arndt, Philipp W. Messer, Dmitri Petrov, Nicolas Galtier, and Laurent Duret. Quantification of gc-biased gene conversion in the human genome. Genome Research, 25(8):1215–1228, 2015.

[19] Cong Guo, Ian C. McDowell, Michael Nodzenski, Denise M. Scholtens, Andrew S. Allen, William L. Lowe, and Timothy E. Reddy. Transversions have larger regulatory effects than transitions. BMC Genomics, 18(1):394, 2017.

[20] Bjarni V. Halldorsson, Marteinn T. Hardarson, Birte Kehr, Unnur Styrkarsdottir, Arnaldur Gylfason, Gudmar Thorleifsson, Florian Zink, Adalbjorg Jonasdottir, Aslaug Jonasdottir, Patrick Sulem, Gisli Masson, Unnur Thorsteinsdottir, Agnar Helgason, Augustine Kong, Daniel F. Gudbjartsson, and Kari Stefansson. The rate of meiotic gene conversion varies by sex and age. Nature Genetics, 48(11):1377, 2016.

[21] Bjarni V. Halldorsson, Gunnar Palsson, Olafur A. Stefansson, Hakon Jonsson, Marteinn T. Hardarson, Hannes P Eggertsson, Bjarni Gunnarsson, Asmundur Oddsson, Gisli H. Halldorsson, Florian Zink, Sigurjon A. Gudjonsson, Michael L. Frigge, Gudmar Thorleifsson, Asgeir Sigurdsson, Simon N. Stacey, Patrick Sulem, Gisli Masson, Agnar Helgason, Daniel F. Gud-b jartsson, Unnur Thorsteinsdottir, and Kari Stefansson. Characterizing mutagenic effects of recombination through a sequence-level genetic map. Science, 363(6425):eaau1043, 2019.

[22] Kelley Harris. Evidence for recent, population-specific evolution of the human mutation rate. Proceedings of the National Academy of Sciences, 112(11):3439–3444, 2015. URL https://www.pnas.org/content/112/11/3439.

[23] Kelley Harris and Jonathan K. Pritchard. Rapid evolution of the human mutation spectrum. eLife, 6:e24284, 2017. URL https://elifesciences.org/articles/24284.

[24] Jude Holmes, Susanna Clark, and Paul Modrich. Strand-specific mismatch correction in nuclear extracts of human and drosophila melanogaster cell lines. Proceedings of the National Academy of Sciences, 87(15):5837–5841, 1990.

[25] Gerald P. Holmquist. Chromosome bands, their chromatin flavors, and their functional features. American Journal of Human Genetics, 51(1):17, 1992. URL https://europepmc.org/article/med/1609794.

[26] Josef Jiricny. The multifaceted mismatch-repair system. Nature Reviews Molecular cell biology, 7(5):335, 2006. URL https://www.nature.com/articles/nrm1907.

[27] Hákon Jónsson, Patrick Sulem, Birte Kehr, Snaedis Kristmundsdottir, Florian Zink, Eirikur Hjartarson, Marteinn T. Hardarson, Kristjan E. Hjorleifsson, Hannes P. Eggertsson, Sigurjon Axel Gudjonsson, Lucas D. Ward, Gudny A. Arnadottir, Einar A. Helgason, Hannes Helgason, Arnaldur Gylfason, Adalbjorg Jonasdottir, Aslaug Jonasdottir, Thorunn Rafnar, Soren Besenbacher, Michael L. Frigge, Simon N. Stacey, Olafur Th. Magnusson, Unnur Thorsteinsdottir, Gisli Masson, Augustine Kong, Bjarni V. Halldorsson, Agnar Helgason, Daniel F. Gudbjartsson, and Kari Stefansson. Whole genome characterization of sequence diversity of 15,220 icelanders. Scientific Data, 4:170115, 2017.

[28] Barbara Kramer, Wilfried Kramer, and Hans-Joachim Fritz. Different base/base mismatches are corrected with different efficiencies by the methyl-directed dna mismatch-repair system of e. coli. Cell, 38(3):879–887, 1984. URL https://www.sciencedirect.com/science/article/pii/0092867484902836.

[29] Heng Li and Richard Durbin. Fast and accurate short read alignment with Burrows–Wheeler transform. Bioinformatics, 25(14):1754–1760, 2009.

[30] Iain Mathieson and David Reich. Differences in the rare variant spectrum among human populations. PLoS Genetics, 13(2):e1006581, 2017. URL https://www.pnas.org/content/112/11/3439.

[31] Aaron McKenna, Matthew Hanna, Eric Banks, Andrey Sivachenko, Kristian Cibulskis, Andrew Kernytsky, Kiran Garimella, David Altshuler, Stacey Gabriel, Mark Daly, and Mark A. DePristo. The Genome Analysis Toolkit: a MapReduce framework for analyzing next-generation DNA sequencing data. Genome Research, 20(9):1297–1303, 2010.

[32] Gilean AT. McVean and Brian Charlesworth. A population genetic model for the evolution of synonymous codon usage: patterns and predictions. Genetics Research, 74:145–158, 1999.

[33] Gilean AT. McVean, Simon R. Myers, Sarah Hunt, Panos Deloukas, David R. Bentley, and Peter Donnelly. The fine-scale structure of recombination rate variation in the human genome. Science, 304(5670):581–584, 2004.

[34] Julien Meunier and Laurent Duret. Recombination drives the evolution of gc-content in the human genome. Molecular Biology and Evolution, 21(6):984–990, 2004. URL https://europepmc.org/article/med/14963104.

[35] Juan I. Montoya-Burgos, Pierre Boursot, and Nicolas Galtier. Recombination explains isochores in mammalian genomes. Trends in Genetics, 19(3):128–130, 2003. URL https://www.sciencedirect.com/science/article/pii/S0168952503000210.

[36] Kasper Munch, Thomas Mailund, Julien Y. Dutheil, and Mikkel Heide Schierup. A fine-scale recombination map of the human-chimpanzee ancestor reveals faster change in humans than in chimpanzees and a strong impact of gc-biased gene conversion. Genome Research, 24(3): 467–474, 2014.

[37] Thomas Nagylaki. Evolution of a large population under gene conversion. Proceedings of the National Academy of Sciences, 80(19):5941–5945, 1983.

[38] Alexander Nater, Maja P. Mattle-Greminger, Anton Nurcahyo, Matthew G. Nowak, Marc De Manuel, Tariq Desai, Colin Groves, Marc Pybus, Tugce Bilgin Sonay, Christian Roos, Adriano R. Lameira, Serge A. Wich, James Askew, Marina Davila-Ross, Gabriella Fredriksson, Guillemde Valles, Ferran Casals, Javier Prado-Martinez, Benoit Goossens, Ernst J. Verschoor, Kristin S. Warren, Ian Singleton, David A. Marques, Joko Pamungkas, Dyah Perwitasari-Farajallah, Puji Rianti, Augustine Tuuga, Ivo G. Gut, Marta Gut, Pablo Orozco-terWengel, Carel P. van Schaik, Jaume Bertranpetit, Maria Anisimova, Aylwyn Scally, Tomas Marques-Bonet, Erik Meijaard, and Michael Krützen. Morphometric, behavioral, and genomic evidence for a new orangutan species. Current Biology, 27(22):3487–3498, 2017.

[39] T Ohta. Slightly deleterious mutant substitutions in evolution. Nature, 246:96–98, 1979.

[40] T Ohta and JH Gillespie. Development of neutral and nearly neutral theories. Theoretical Population Biology, 49(2):128–142, 1996.

[41] Eugénie Pessia, Alexandra Popa, Sylvain Mousset, Clément Rezvoy, Laurent Duret, and Gabriel AB. Marais. Evidence for widespread gc-biased gene conversion in eukaryotes. Genome Biology and Evolution, 4(7):675–682, 2012. URL https://academic.oup.com/gbe/article/4/7/675/740757.

[42] Javier Prado-Martinez, Peter H. Sudmant, Jeffrey M. Kidd, Heng Li, Joanna L. Kelley, Belen Lorente-Galdos, Krishna R. Veeramah, August E. Woerner, Timothy D. O’connor, Gabriel Santpere, Alexander Cagan, Christoph Theunert, Ferran Casals, Hafid Laayouni, Kasper Munch, Asger Hobolth, Anders E. Halager, Maika Malig, Jessica Hernandez-Rodriguez, Irene Hernando-Herraez, Kay Prüfer, Marc Pybus, Laurel Johnstone, Michael Lachmann, Can Alkan, Dorina Twigg, Natalia Petit, Carl Baker, Fereydoun Hormozdiari, Marcos Fernandez-Callejo, Marc Dabad, Michael L. Wilson, Laurie Stevison, Cristina Camprubí, Tiago Carvalho, Aurora Ruiz-Herrera, Laura Vives, Marta Mele, Teresa Abello, Ivanela Kondova, Ronald E. Bontrop, Anne Pusey, Felix Lankester, John A. Kiyang, Richard A. Bergl, Elizabeth Lonsdorf, Simon Myers, Mario Ventura, Pascal Gagneux, David Comas, Hans Siegismund, Julie Blanc, Lidia Agueda-Calpena, Marta Gut, Lucinda Fulton, Sarah A. Tishkoff, James C. Mullikin, Richard K. Wilson, Ivo G. Gut, Mary Katherine Gonder, Oliver A. Ryder, Beatrice H. Hahn, Arcadi Navarro, Joshua M. Akey, Jaume Bertranpetit, David Reich, Thomas Mailund, Mikkel H. Schierup, Christina Hvilsom, Aida M. Andrés, Jeffrey D. Wall, Carlos D. Bustamante, Michael F. Hammer, Evan E. Eichler, and Tomas Marques-Bonet. Great ape genetic diversity and population history. Nature, 499(7459):471, 2013.

[43] Jonathan Romiguier, Vincent Ranwez, Emmanuel JP. Douzery, and Nicolas Galtier. Contrasting gc-content dynamics across 33 mammalian genomes: relationship with lifehistory traits and chromosome sizes. Genome research, 20(8):1001–1009, 2010. URL https://genome.cshlp.org/content/20/8/1001.

[44] P. Savatier, G. Trabuchet, C. Faure, Y. Chebloune, Manolo Gouy, Ghislain Verdier, and VM. Nigon. Evolution of the primate β-globin gene region: High rate of variation in cpg dinucleotides and in short repeated sequences between man and chimpanzee. Journal of Molecular Biology, 182(1):21–29, 1985.

[45] Paul Scheet and Matthew Stephens. A fast and flexible statistical model for large-scale population genotype data: applications to inferring missing genotypes and haplotypic phase. The American Journal of Human Genetics, 78(4):629–644, 2006.

[46] Dominik Schrempf and Asger Hobolth. An alternative derivation of the stationary distribution of the multivariate neutral wright–fisher model for low mutation rates with a view to mutation rate estimation from site frequency data. Theoretical Population Biology, 114:88–94, 2017.

[47] Eric U. Selker and Judith N. Stevens. Dna methylation at asymmetric sites is associated with numerous transition mutations. Proceedings of the National Academy of Sciences, 82(23):8114–8118, 1985.

[48] Vladimir B. Seplyarskiy, Peter Kharchenko, Alexey S. Kondrashov, and Georgii A. Bazykin. Heterogeneity of the transition/transversion ratio in drosophila and hominidae genomes. Molecular Biology and Evolution, 29(8):1943–1955, 2012.

[49] AFA Smit, R Hubley, and P Green. Repeatmasker open-3.0. Institute for Systems Biology, 2004. URL http://www.repeatmasker.org.

[50] Jeffrey P. Spence and Yun S. Song. Inference and analysis of population-specific fine-scale recombination maps across 26 diverse human populations. S’cience Advances, 5(10):eaaw9206, 2019.

[51] Maria Spies and Richard Fishel. Mismatch repair during homologous and homeologous recombination. Cold Spring Harbor Perspectives in Biology, 7(3):a022657, 2015. URL https://cshperspectives.cshlp.org/content/7/3/a022657.

[52] Matthew Stephens, Nicholas J. Smith, and Peter Donnelly. A new statistical method for haplotype reconstruction from population data. The American Journal of Human Genetics, 68(4): 978–989, 2001.

[53] Laurie S. Stevison, August E. Woerner, Jeffrey M. Kidd, Joanna L. Kelley, Krishna R. Veeramah, Kimberly F. McManus, Great Ape Genome Project, Carlos D. Bustamante, Michael F. Hammer, and Jeffrey D. Wall. The time scale of recombination rate evolution in great apes. Molecular Biology and Evolution, 33(4):928–945, 2015.

[54] Shin-San Su, Robert S. Lahue, Karin G. Au, and Paul Modrich. Mispair specificity of methyl-directed dna mismatch correction in vitro. Journal of Biological Chemistry, 263(14):6829–6835, 1988. URL https://www.jbc.org/content/263/14/6829.

[55] Artem Tarasov, Albert J. Vilella, Edwin Cuppen, Isaac J. Nijman, and Pjotr Prins. Sambamba: fast processing of NGS alignment formats. Bioinformatics, 31(12):2032–2034, 2015.

[56] David C. Thomas, JD. Roberts, and TA. Kunkel. Heteroduplex repair in extracts of human hela cells. Journal of Biological Chemistry, 266(6):3744–3751, 1991.

[57] Flavie Tortereau, Bertrand Servin, Laurent Frantz, Hendrik-Jan Megens, Denis Milan, Gary Rohrer, Ralph Wiedmann, Jonathan Beever, Alan L Archibald, Lawrence B. Schook, and Martien AM. Groenen. A high density recombination map of the pig reveals a correlation between sex-specific recombination and gc content. BMC Genomics, 13(1):586, 2012. URL https://bmcgenomics.biomedcentral.com/articles/10.1186/1471-2164-13-586.

[58] Claus Vogl and Juraj Bergman. Inference of directional selection and mutation parameters assuming equilibrium. Theoretical Population Biology, 106:71–82, 2015.

[59] Claus Vogl and Florian Clemente. The allele-frequency spectrum in a decoupled moran model with mutation, drift, and directional selection, assuming small mutation rates. Theoretical Population Biology, 81(3):197–209, 2012.

[60] Karin Wiebauer and Josef Jiricny. In vitro correction of g o t mispairs to g o c pairs in nuclear extracts from human cells. Nature, 339(6221):234, 1989.

[61] Karin Wiebauer and Josef Jiricny. Mismatch-specific thymine dna glycosylase and dna polymerase beta mediate the correction of gt mispairs in nuclear extracts from human cells. Proceedings of the National Academy of Sciences, 87(15):5842–5845, 1990.

[62] Amy L. Williams, Giulio Genovese, Thomas Dyer, Nicolas Altemose, Katherine Truax, Goo Jun, Nick Patterson, Simon R. Myers, Joanne E. Curran, Ravi Duggirala, John Blangero, David Reich, and Molly Przeworski on behalf of the T2D-GENES Consortium. Non-crossover gene conversions show strong gc bias and unexpected clustering in humans. eLife, 4:e04637, 2015.

[63] Yali Xue, Javier Prado-Martinez, Peter H. Sudmant, Vagheesh Narasimhan, Qasim Ayub, Michal Szpak, Peter Frandsen, Yuan Chen, Bryndis Yngvadottir, David N. Cooper, Marc de Manuel, Jessica Hernandez-Rodriguez, Irene Lobon, Hans R. Siegismund, Luca Pagani, Michael A. Quaill, Christina Hvilsom, Antoine Mudakikwa, Evan E. Eichler, Michael R. Cranfield, Tomas Marques-Bonet, Chris Tyler-Smith, and Aylwyn Scally. Mountain gorilla genomes reveal the impact of long-term population decline and inbreeding. Science, 348(6231):242–245, 2015.

[64] Jianzhi Zhang. Rates of conservative and radical nonsynonymous nucleotide substitutions in mammalian nuclear genes. Journal of Molecular Evolution, 50(1):56–68, 2000.

